# MEMO1 is a Metal Containing Regulator of Iron Homeostasis in Cancer Cells

**DOI:** 10.1101/2023.02.28.530460

**Authors:** Natalia Dolgova, Eva-Maria E. Uhlemann, Michal T. Boniecki, Frederick S. Vizeacoumar, Martina Ralle, Marco Tonelli, Syed A. Abbas, Jaala Patry, Hussain Elhasasna, Andrew Freywald, Franco J. Vizeacoumar, Oleg Y. Dmitriev

## Abstract

Mediator of ERBB2-driven Cell Motility 1 (MEMO1) is an evolutionary conserved protein implicated in many biological processes; however, its primary molecular function remains unknown. Importantly, MEMO1 is overexpressed in many types of cancer and was shown to modulate breast cancer metastasis through altered cell motility.

To better understand the function of MEMO1 in cancer cells, we analyzed genetic interactions of MEMO1 using gene essentiality data from 1028 cancer cell lines and found multiple iron-related genes exhibiting genetic relationships with MEMO1. We experimentally confirmed several interactions between MEMO1 and iron-related proteins in living cells or *in vitro*, most notably, the iron transporters transferrin (*TF*), transferrin receptor 2 (*TFR*2), and mitoferrin-2 (*SLC25A28*), and the global iron response regulator IRP1 (*ACO1*). These interactions indicate that cells with high MEMO1 expression levels are hypersensitive to the disruptions in iron distribution. Our data also indicate that MEMO1 is involved in ferroptosis and is linked to iron supply to mitochondria.

We have found that purified MEMO1 binds iron with high affinity under redox conditions mimicking intracellular environment and solved MEMO1 structures in complex with iron and copper. Our work reveals that the iron coordination mode in MEMO1 is very similar to that of iron-containing extradiol dioxygenases, which also display a similar structural fold. We conclude that MEMO1 is an iron-binding protein that regulates iron homeostasis in cancer cells.

## Introduction

MEMO1 is a highly conserved protein found in the cytosol of eukaryotic cells from yeast to humans. MEMO1 appears to play a crucial role in cell motility, and has been linked to several biological processes, but its primary function remains unknown. MEMO1 has been implicated in lifespan changes in *C. elegans* and mice (1, 2), regulation of vitamin D metabolism (3), and bone and central nervous system development (4–6).

In cancer context, MEMO1 supports the ability of breast tumor cells to invade surrounding tissues, leading to metastasis (7–9). Knockdown of MEMO1 expression reduces breast cancer cell migration in culture, and significantly suppresses lung metastasis in a xenograft model (7). In the clinical setting, retrospective analysis of resected tumors showed a strong correlation between increased expression of MEMO1 and reduced patient survival (7). These effects have been linked to the interaction between the ERBB2 (HER2) receptor and MEMO1, which in turn was proposed to relay the activation of ERBB receptor heterodimers to the microtubule cytoskeleton, thus inducing growth of lamellipodia and enabling cancer cell migration (8). The interaction with ERBB2 gave MEMO1 its name (**M**ediator of **E**RBB2-driven Cell **Mo**tility **1**). MEMO1 also contributes to breast carcinogenesis through the insulin receptor substrate protein 1 (IRS1) pathway (10) and through the interaction with the extranuclear estrogen receptor (11, 12).

MEMO1 was shown to catalyze copper-dependent redox reactions, such as superoxide radical production (7). This led to the idea that MEMO1 is required for sustained reactive oxygen species (ROS) production, likely in conjunction with NADPH-oxidase 1 (NOX1) activation. ROS are known to modulate functions of several proteins required for cell motility (13). Alternatively, MEMO1 has been proposed to protect cells from ROS generation by sequestering copper (14). At the same time, sequence homology and structure analyses revealed strong similarity of MEMO1 to iron-containing dioxygenases, redox enzymes that catalyze the incorporation of molecular oxygen into organic molecules (15), suggesting that MEMO1 may have other, iron-dependent, functions in the cell. Strikingly, within the very distinct cluster of four top matches to MEMO1 structure found in the Protein Data Bank using DALI (16), three proved to be iron dioxygenases, with the fourth being a putative dioxygenase.

Additional clues to MEMO1 function were offered by the genome wide studies of genetic interactions in yeast. *MEMO1* homolog in *S. cerevisiae* is *MHO1*, with 37% amino acid residue identity. In the global network of genetic interactions in yeast (17), the highest interaction profile similarity was found between *MHO1* and *LSO1*, which encodes a protein strongly induced in response to iron deprivation, and a likely component of the iron transport pathway in yeast (18). Among the five highest scoring hits in the profile similarity search, there was another iron linked protein, the ferredoxin reductase ARH1, an iron-sulfur protein playing an essential role in [2Fe-2S] cluster biogenesis (19). Thus, iron mediated redox reactions emerged as a common MEMO1 denominator from two orthogonal bioinformatics approaches, prompting us to investigate its iron connection.

In the present work, we show that MEMO1 is an iron-binding protein involved in iron metabolism in the cell. Iron has been implicated in carcinogenesis, tumor growth, and metastasis, in particular in breast cancer, by multiple lines of evidence, due to its potentially disruptive role in redox balance in the cell and also because of elevated iron requirements of the rapidly proliferating cancer cells (20, 21). Thus, MEMO1 emerges as a direct molecular link between iron metabolism and metastasis in breast cancer.

## Results

### MEMO1 displays genetic interactions with multiple iron related proteins

Our analysis of the Cancer Genome Atlas (https://www.cancer.gov/tcga) data reveals that, in addition to breast cancer (*P<*10^-29^ by Mann-Whitney U-test), MEMO1 is overexpressed in the malignancies of colon, lung and uterine origins (*P<*0.0001), among others, while kidney, head and neck squamous cell tumors and melanoma show little or no difference in median expression levels (Fig. 1A and 1B). Further analyses of MEMO1 levels in breast cancer subtypes determined that MEMO1 overexpression is more prominent in triple negative (TNBC) and HER2+ (HER2 enriched) breast tumors than in the luminal subtype (Fig. 1C). This makes MEMO1 a potential therapeutic target in HER2+ breast cancer and, importantly, in TNBC, where few effective treatment options exist, and patient survival is poor. Therefore, we set out to investigate the molecular function of MEMO1.

**Fig. 1.**
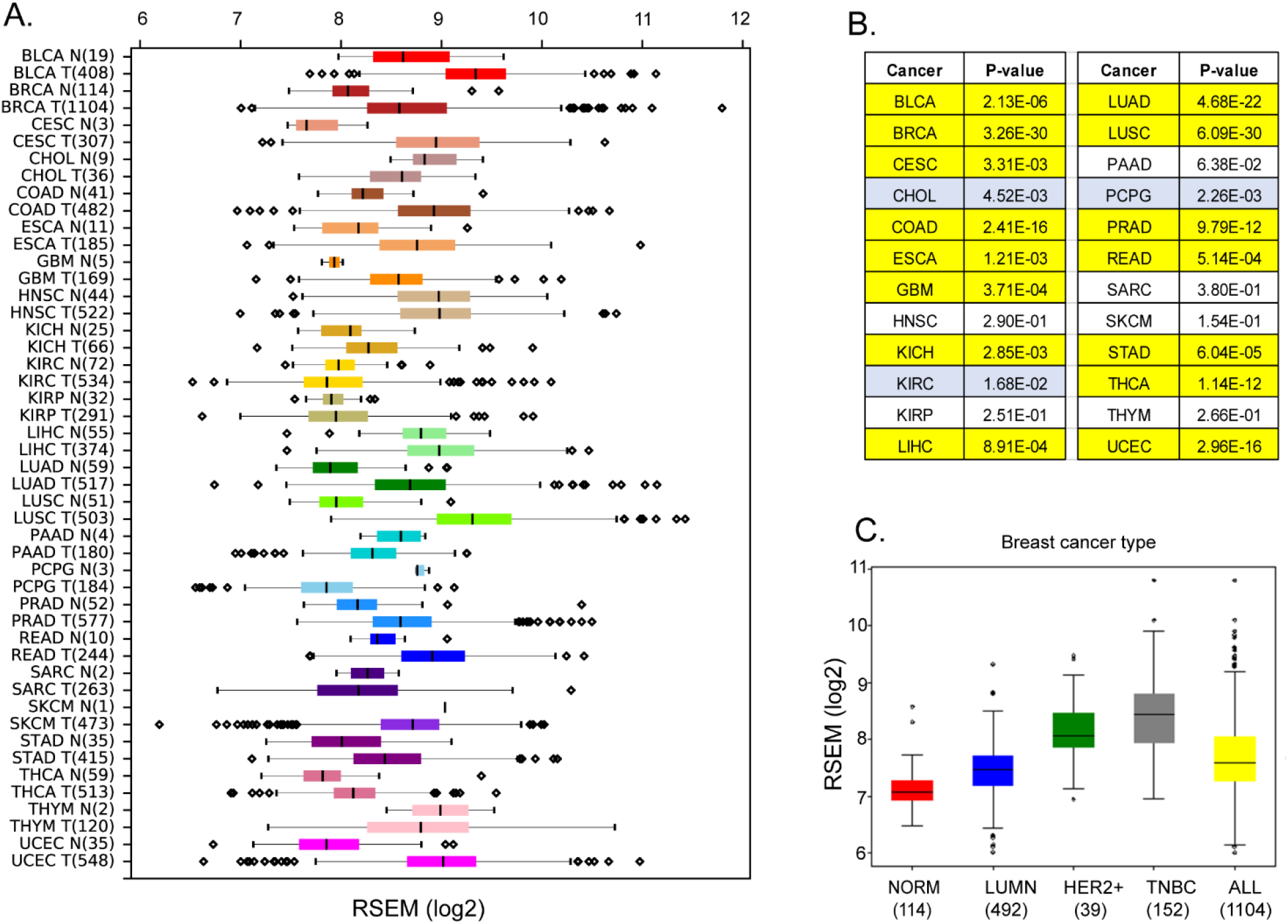
Expression levels of MEMO1 in tumors and corresponding normal tissue in various malignancies. The data is from the Cancer Genome Atlas (TCGA) database (https://www.cancer.gov/tcga) **(A)** Tissue profile of MEMO1 expression in cancer. Standard TCGA cancer type abbreviations are used (BRCA – breast cancer, SKCM – melanoma). **(B)** Statistically significant differences in MEMO1 expression levels between the tumors and the corresponding normal tissue (*P<*0.05 by Mann-Whitney U-test) are highlighted in *yellow* (higher expression in tumors) or *blue* (lower expression in tumors). **(C)** Subtype analysis of MEMO1 expression in breast cancer, including luminal (*blue*), HER2 positive (*green*) and triple-negative breast cancer (TNBC, *grey*). All differences vs. normal breast tissue (red) are highly significant (*P<*10^-17^). Note the logarithmic scale (RSEM log_2_) on the transcript level axis in A and C.

Structural similarity of MEMO1 to the iron containing dioxygenases and a clear link between MEMO1 homolog MHO1 and iron metabolism that emerged from genome wide analyses of genetic interactions networks in yeast led us to a hypothesis that MEMO1 plays a role in iron metabolism in cancer cells. As genetic interactions (GIs) are known to be functionally coherent (22), we applied a novel approach to identify GIs of MEMO1 using publicly available gene essentiality data from multiple cell lines. We called a given gene to exhibit GI with MEMO1 if this gene was found to become essential only in cancer cells that either overexpress or downregulate MEMO1. Using previously published genome-wide screens, we analyzed gene essentiality scores derived from 1028 cancer cell lines (23–29) displaying a wide range of MEMO1 expression levels.

This analysis yielded genes that become highly essential (Wilcoxon-rank sum test *P*<0.05), when MEMO1 is either under-expressed, representing GIs identified from loss of function (LOF-GIs), or overexpressed, representing GIs identified from gain of function, (GOF-GIs). Since most of the previous work on MEMO1 and its role in cancer was done in breast cancer cell lines, we analyzed the breast cancer and pan-cancer datasets separately.

Consistent with the known roles of MEMO1 (30), we found its GIs to be enriched in the following gene sets: Cell Adhesion Molecule Binding (False Discovery Rate, FDR 1.13E-03), Response To Insulin (FDR 2.07E-03), Insulin Signaling Pathway (FDR 6.83E-03), Cellular response to chemical stress (FDR 2.05E-04), ERBB Signaling Pathway (FDR 3.47E-04), and Focal Adhesion (FDR 1.99E-02) (Fig. S1) for LOF-GIs, while gene set enrichment analysis (GSEA) of GOF-GIs identified Signaling by ERBB2 (FDR < 1.00E-05), Microtubule Cytoskeleton Organization (FDR 8.29E-04), Extra-nuclear Estrogen Signaling (FDR 6.13E-04) (Fig. S2). Strong enrichment of the gene sets directly related to the known MEMO1 functions convincingly validated our approach.

Remarkably, gene set enrichment analysis (GSEA) of the GOF-GIs identified multiple partially overlapping gene sets related to the mitochondrial energy metabolism and redox balance in the cell, including Mitochondrial Transport (FDR, 5.64E-04), Mitochondrial Electron Transport NADH to Ubiquinone (FDR 8.92E-04), Respiratory Electron Transport Chain (FDR 1.54E-02), Citric Acid (TCA) Cycle and Respiratory Electron Transport (FDR 5.99E-03), Oxidative Phosphorylation (FDR 4.32E-02), Glutathione Metabolism (FDR 1.45E-01) and Oxidative Stress Induced Senescence (FDR 1.70E-02) (Fig. S2). These enrichment categories include multiple genes encoding iron containing proteins, as well as those involved in iron-dependent cellular processes, such as ferroptosis. Following this lead, we asked if enrichment of these gene sets in the MEMO1 network of the GOF-GIs may reflect specifically high sensitivity of high-MEMO1 cancer cells to the disruptions of iron metabolism.

Further analysis identified sixteen genes encoding proteins involved in iron metabolism and iron transport that showed statistically significant GIs with *MEMO1* (Tables S1 and S2, Fig. S3 and S4). Ten genes were found to be more essential in cancer cell lines with high expression of MEMO1 (GOF-GIs, Table S1), while five genes were more essential in the cell lines with low MEMO1 levels (LOF-GIs, Table S2). For example, knockout or knockdown of transferrin receptor 2 (TFR2), iron transporter mitoferrin-2 (SLC25A28), or iron response protein (ACO1) selectively suppresses proliferation of the high-MEMO1 cell lines, while knockout of iron transporter DMT1 (SLC11A2) leads to a stronger inhibition of proliferation in the cell lines with low MEMO1 expression. The other iron related genes showing genetic interactions with *MEMO1* encode mitochondrial proteins that are involved in iron-sulfur cluster biogenesis (*HSPA9, ISCU*, *FXN, ISCA2, NUBPL*), contain iron-sulfur clusters (*ACO2, LIAS*), or are involved in heme synthesis (*HMOX1*). As these results strongly support a link between MEMO1 and iron related proteins, we next explored several of these interactions in more detail.

### MEMO1 regulates iron homeostasis in the cell

To confirm the functional link between iron homeostasis and MEMO1 experimentally, we generated clonal MEMO1 knockdowns and knockouts in MDA-MB-231 triple negative breast cancer and A-375 melanoma cell lines using the CRISPR-Cas9 technology and compared the effects of shRNA knockdown of selected genes involved in iron homeostasis on the proliferation rates of the cells with high (parental), low (knockdown) and no (knockout) MEMO1 expression. We chose a melanoma cell line for comparison, because, like breast cancer, it showed a high level of MEMO1 expression, but, unlike breast cancer, there is no statistically significant difference between MEMO1 expression levels in melanoma and normal skin tissue (Fig. 1B). Thus, melanoma serves as a useful benchmark for a cancer, where MEMO1 overexpression is not required to support malignant transformation. In agreement with the previous reports (7, 8), we found that MEMO1 knockout in breast cancer cells results in the loss of cell motility as assessed by the wound healing assay (Fig. S5A). MEMO1 knockdown and knockout also decreased overall rates of breast cancer cell proliferation (Fig. S5B). As expected, no strong motility or proliferation rate dependence on MEMO1 expression level was observed in the melanoma cells (Figs. S5C and S5 D).

Because MEMO1 GIs identified by the gene essentiality analysis from 1028 cell lines clearly suggested a link between MEMO1 and iron transport, we started by investigating the effect of suppressing transferrin receptors TFR1 and TFR2. TFR1 is the key receptor essential for iron uptake, while TFR2 appears to be involved in the regulation of iron homeostasis rather than performing bulk iron uptake (31). TFR2, but not TFR1, was found to exhibit GI with MEMO1 along with other iron-related genes (Fig. 2A). The shRNA targeting *TFR1* decreased TFR1 protein expression by more than 90% and caused near complete proliferation inhibition in all the tested cell lines regardless of MEMO1 expression level (Fig. 2B, S6A and S6B). Thus, TFR1 appears to be an essential protein as its loss is nearly lethal on its own, explaining why *TFR1* was not detected in our *in silico* screening of MEMO1 genetic interactions.

**Fig. 2.**
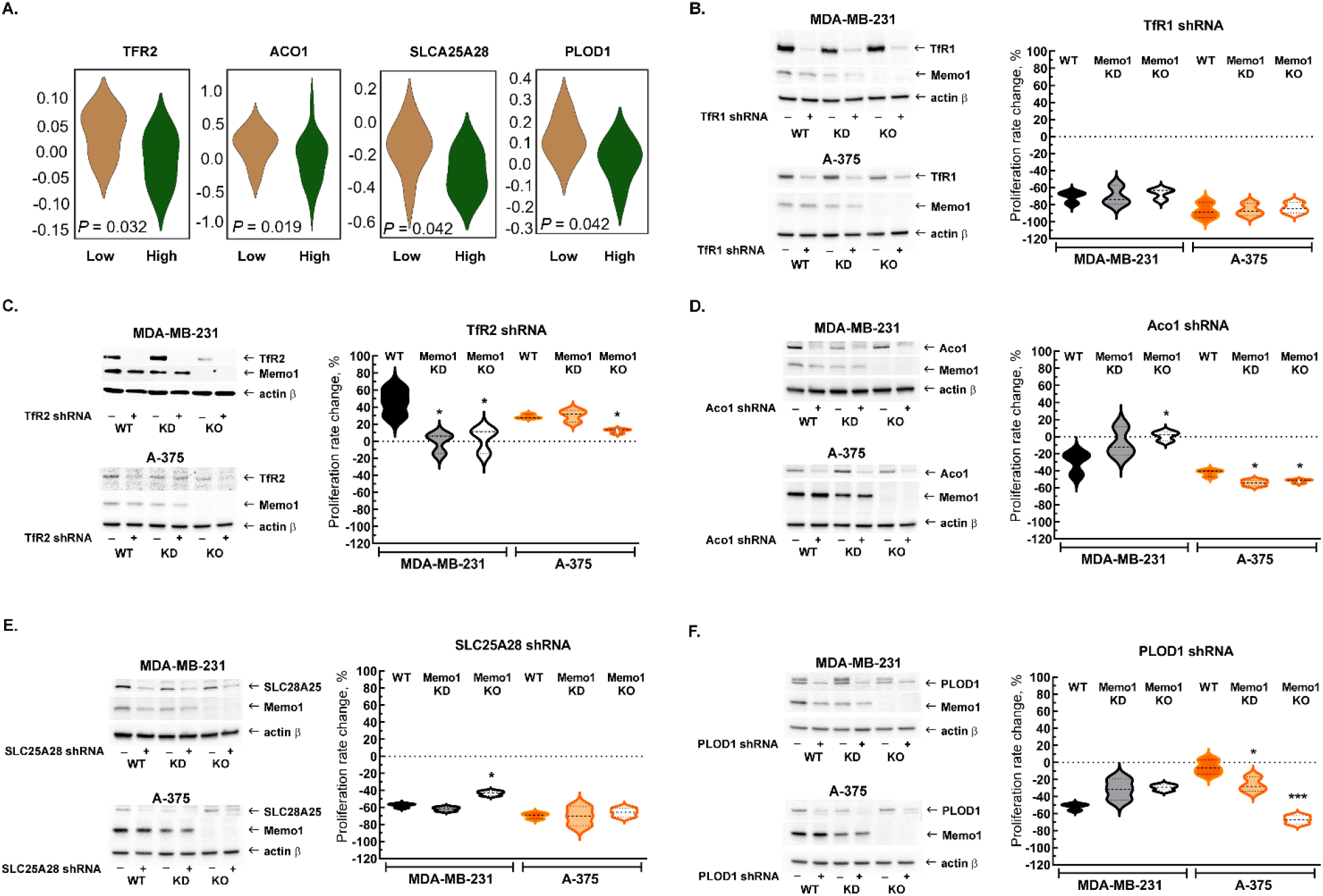
Interactions between *MEMO1* and iron-related genes. **(A)** Gene essentiality score distribution for the selected genes in the high- and low-MEMO1 expressing groups in multiple cancer cell lines as shown by the database analysis (cf. Table S1). Effects of the TFR1 **(B)**, TFR2 **(C)**, ACO1 **(D)**, SLC25A28 **(E)** and PLOD1 **(F)** shRNA knockdowns at different MEMO1 expression levels (WT – parental cell line, KD – *MEMO1* knockdown, KO – *MEMO1* knockout). Protein levels were detected by Western blot (*left panels*). The relative shRNA effect on cell proliferation (*right panels*) is expressed as the difference between the proliferation rates with the shRNA targeting the gene of interest and the control shRNA (against the red fluorescent protein) divided by the proliferation rate with the control shRNA.

In contrast to *TFR1*, downregulation of TFR2 by shRNA in the high-MEMO1 breast cancer cell line results in the *activation* of cell proliferation, possibly due to the increased availability of iron. By comparison, breast cancer cells with MEMO1 knockout or knockdown show no proliferation rate change in response to TFR2 shRNA knockdown (Fig. 2C and S6C), indicating that TFR2 has a GOF-GI with MEMO1. A similar MEMO1-dependent growth activation pattern was also observed in melanoma cell lines (Fig. 2C and S6D).

We have also experimentally tested GIs between *MEMO1* and several other iron-related genes detected in our *in silico* analyses (Fig. 2A) and chosen for their key role in the regulation of iron homeostasis in the cell (*ACO1*), iron transport and processing in mitochondria (*SLC25A28* and *HSPA9*), or a direct link between the known MEMO1 function in modulating cancer cell motility and iron (*PLOD1*). *ACO1, SLC25A28* and *PLOD1* showed GOF interactions with MEMO1 in breast cancer cells. Remarkably, *ACO1* knockdown only inhibited proliferation of the high-MEMO1 cells, but not MEMO1 knockout breast cancer cells (Fig. 2D), while *SLC25A28* (Fig. 2E and S7A) and *PLOD1* (Fig. 2F and S8A) knockdowns showed some effect in all cell lines, but a stronger inhibition in the high-MEMO1 cells than in MEMO1 knockout. By comparison, none of the three genes showed GOF interactions in melanoma cells (Fig. 2D, 2E, 2F, S7B and S8B). In fact, *ACO1* and *PLOD1* displayed LOF interactions in melanoma. Like TFR1, shRNA knockdown of HSPA9 results in a major proliferation inhibition of breast cancer and, particularly, of melanoma cells, regardless of MEMO1 expression level (Fig. S9).

Next, we investigated correlations between the expression levels of MEMO1 and the six iron related proteins whose genetic interactions with MEMO1 had been confirmed. Analysis of the gene expression levels in multiple breast cancer cell lines (Fig. 3A, Fig. S10A) revealed weak but statistically highly significant correlations between the levels of MEMO1 and TFR1 (*P=*4.6E-18), TFR2 (*P=*1.8E-14) and PLOD1 (*P=*1.8E-33). By comparison, none of the correlations in melanoma exceeded Spearman’s rank correlation coefficient of 0.2 (Fig. 3B, Fig. S10), a common, if arbitrary, correlation detection threshold.

**Fig. 3.**
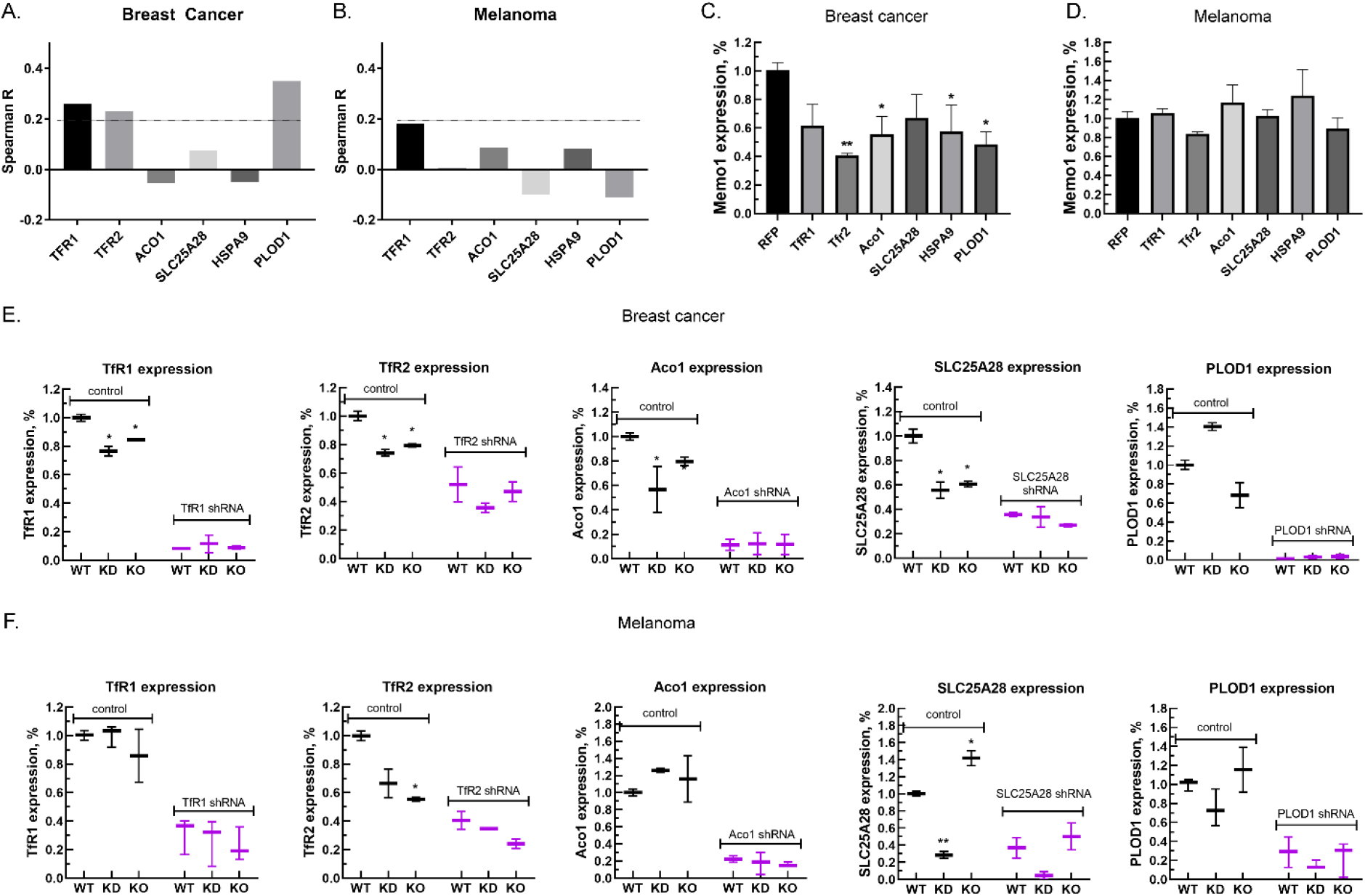
Relationship between the expression levels of MEMO1 and iron-related proteins in breast cancer and melanoma. **(A, B)** Correlations between the expression levels of MEMO1 and selected iron related proteins in multiple breast cancer **(A)** and melanoma **(B)** cell lines (cf. Fig. S10) as measured by Spearman rank-order correlation coefficient. **(C, D)** Effect of the selected iron-related shRNA gene knockdowns on MEMO1 levels in breast cancer **(C)** and melanoma **(D)** cell lines detected by Western blot analysis (cf. Fig. 2B - 2F). **(E, F)** Levels of the selected iron-related proteins at various MEMO1 expression levels in breast cancer **(E)** and melanoma **(F)** detected by Western blot analysis in control (shRNA against the red fluorescent protein) and with the shRNA targeting the gene of interest.

Experimentally, knocking down TFR1, TFR2, ACO1, HSPA9, and PLOD1 in the breast cancer cell line MDA-MB-231 resulted in a decrease in the expression level of MEMO1 by 40-60% (Fig. 3C), but, as expected, no significant effect in the melanoma cell line A-375 was observed (Fig. 3D). Conversely, knockdown or knockout of MEMO1 in breast cancer cells resulted in a statistically significant decrease in the expression levels of TFR1, TFR2, ACO1, and SLC25A28 (Fig. 3E). By comparison, MEMO1 knockdown or knockout in melanoma cells decreased only the expression level of TFR2 (Fig. 3F), the effect on SLC25A28 knockdown being difficult to interpret. Thus, expression levels of MEMO1 and the other iron-related proteins are correlated in TNBC, suggesting regulation through a common mechanism.

### MEMO1 plays an important role in the maintenance of iron concentration in mitochondria

Genetic interactions and expression level correlations between MEMO1 and the iron transport and iron regulating proteins suggested that MEMO1 may also be involved in regulating iron levels in the cell. To test this hypothesis, we measured iron concentration in the cytosol and in the crude mitochondrial fractions of MDA-MB-231 cells in comparison to the same fractions of MEMO1 knockdown and knockout cell lines (M67-2 and M67-9). Cells with MEMO1 knockout showed significantly lower iron concentrations in both the cytosolic (Fig. 4A) and the mitochondrial (Fig. 4B) fractions. These results suggest that overexpression of MEMO1 allows cells to accumulate higher levels of iron, needed for maintaining the accelerated cell proliferation. This effect may be mediated by a TFR2 - MEMO1 pathway through the iron transport protein transferrin, consistent both with the genetic interactions between MEMO1 and TFR2 and with the physical interaction between the purified MEMO1 and transferrin observed in our experiments by microscale thermophoresis (Fig. 4C). Interestingly, although both *apo-* and *holo-*transferrin bound MEMO1, the *apo-*form showed approximately tenfold higher affinity for MEMO1 (*K_d_=*7.9 × 10^-9^ M) compared to the *holo*-form (*K_d_=*9.7 × 10^-8^ M).

**Fig. 4.**
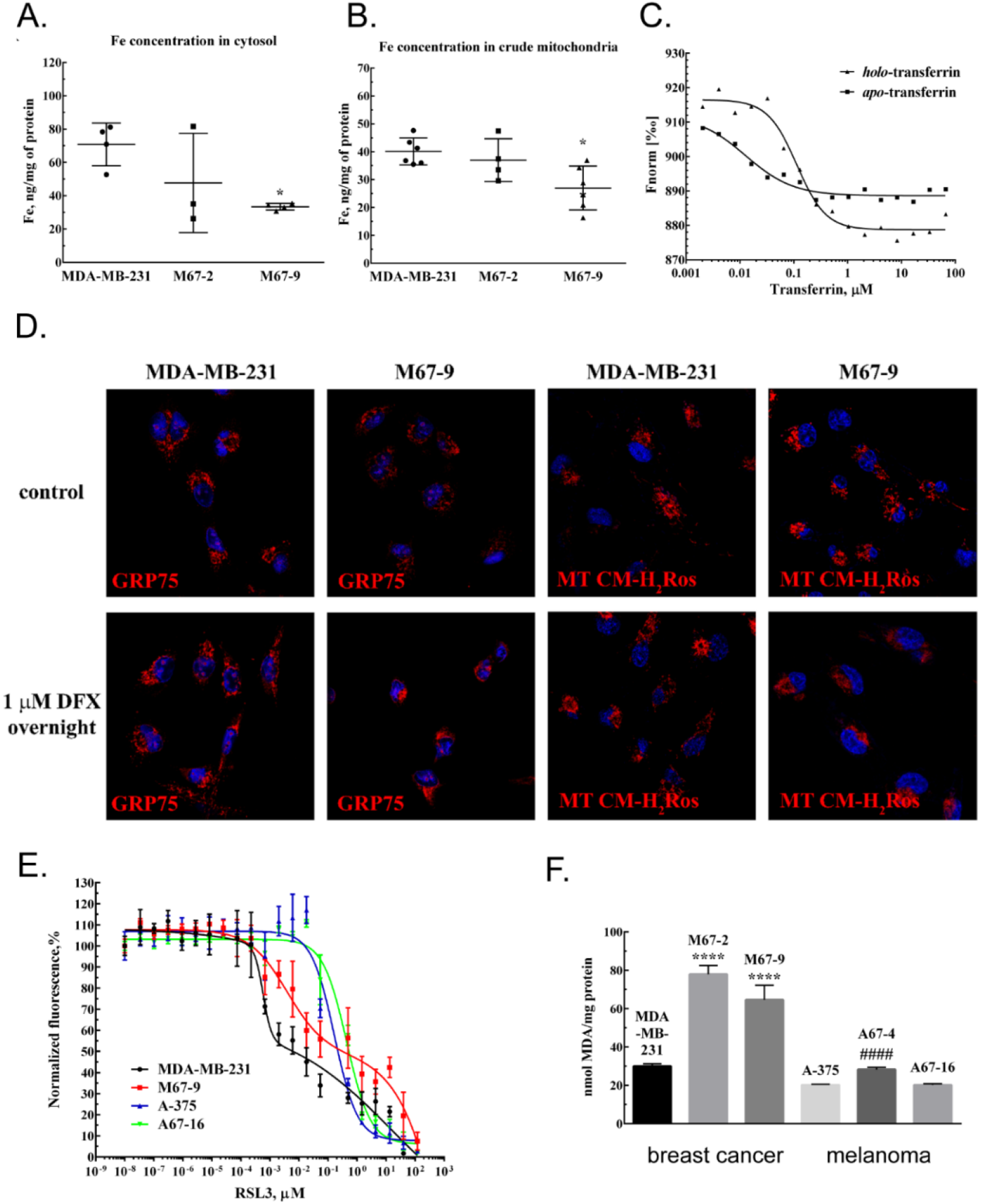
MEMO1 expression affects iron levels, mitochondrial morphology, and ferroptosis sensitivity of the cells. **(A, B)** Iron levels in the cytosolic **(A)** and mitochondrial **(B)** fractions from breast cancer cells with high (MDA-MB-231), low (M67-2), and no (M67-9) MEMO1 expression. **(C)** MEMO1 binding to *apo-* and *holo-*transferrin, as measured by the infrared laser-induced thermophoresis. **(D)** MEMO1 knockout in breast cancer cells M67-9 results in perinuclear mitochondrial clustering in the presence of iron chelator deferoxamine (DFX). Mitochondrial marker Grp75 (HSPA9) and MitoTracker CM-H_2_Ros are *red*, DAPI-stained nuclei are *blue*. **(E)** MEMO1 knockout in breast cancer (M67-9) increases resistance to the ferroptosis inducer RSL3, compared to the parental cell line MDA-MB-231, with an insignificant knockout effect in melanoma (A67-16 and A-375 respectively). **(F)** Malondialdehyde assay shows lower rates of lipid oxidation in the breast cancer cells with high MEMO1 expression (MDA-MB-231) compared to cells with MEMO1 knockout (M67-9) and knockdown (M67-2) cells. The difference is smaller in melanoma cells (A67-4 is MEMO1 knockdown and A67-16 is the knockout).

The decrease in mitochondrial iron in MEMO1-knockout cells taken together with the interactions between MEMO1 and the mitochondrial iron transporter SLC25A28 (mitoferrin-2) prompted us to investigate how MEMO1 knockout affects mitochondria. Mitochondrial morphology in MDA-MB-231 cells with and without MEMO1 knockout was evaluated using mitochondrial marker GRP75 (HSPA9) and the transmembrane electric potential-sensitive dye MitoTracker CM-H2Ros (Fig. 4D). Under basal culture conditions, both the wild type and MEMO1-knockout cell lines display normal mitochondrial shape. By comparison, cells incubated overnight with 1 µM iron chelator deferoxamine (DFX) present normal distribution of mitochondria in the wild type cells, while cells with MEMO1 knockout (M67-9) show perinuclear mitochondrial clustering, as detected both by GRP75 distribution and MitoTracker CM-H_2_Ros staining.

This observation indicates that depletion of labile iron pool caused by the incubation with DFX results in changes in mitochondrial morphology, but only in the cells without MEMO1. Thus, MEMO1 not only plays an important role in overall iron homeostasis in the cell but is specifically involved in iron regulation in mitochondria.

### MEMO1 promotes ferroptosis via increase in iron concentration in the cell

As described above, we found that MEMO1 is involved in the regulation of iron concentration in the cells. Elevated iron levels are known to make cells more susceptible to ferroptosis, a type of programmed cell death triggered by the iron-dependent lipid oxidation, and distinct from apoptosis, necrosis, autophagy, and other types of cell death (32). Further suggesting a possible involvement of MEMO1 in ferroptosis, we found GOF-GI between MEMO1 and most of the proteins involved in mitochondria-based ferroptosis (33), including aconitase, VDACs, mitoferrin-2 (SLC25A28), frataxin (FXN), citrate synthase (CS), Acyl-CoA synthetase long-chain family member 4 (ACSL4), sterol carrier protein 2 (SCP2), sirtuin 3 (SIRT3), glutaminase 2 (GLS2), SCO2, and fumarate hydratase (FH) (Supplementary Table S3). To probe the role of MEMO1 in ferroptosis, we compared responses of the wild type and MEMO1-knockout MDA-MB-231 and A-375 cells to the ferroptosis-inducing agent RSL3 (32), a specific inhibitor of glutathione peroxidase 4 (GPX4), the enzyme that resides in mitochondria and plays an essential role in protecting cells against lipid oxidation (34).

MEMO1 knockout results in a decreased cytotoxicity of RSL3 in breast cancer cells (Fig. 4E), suggesting that high-MEMO1 cells are more sensitive to ferroptosis. Lipid oxidation is one of the hallmarks of ferroptosis. Breast cancer cells with MEMO1 knockdown and knockout have significantly higher levels of lipid oxidation product malondialdehyde (MDA) (Fig. 4F). Taken together, these results indicate that MEMO1 knockout increases ferroptosis resistance in breast cancer cells through the reduced labile iron pool (Fig. 4A and 4B) and a higher tolerance towards lipid oxidation products. Melanoma cells show much smaller MEMO1-dependent variations in the MDA level than breast cancer cells, consistent with the observed smaller difference in RSL3 sensitivity between the wild type and MEMO1 knockout in melanoma (Fig. 4E). Overall, these results indicate that MEMO1 overexpression sensitizes breast cancer cells to ferroptosis.

### MEMO1 is an iron-binding protein

We have discovered GIs between MEMO1 and many iron-dependent proteins and found strong indications of MEMO1 involvement in the regulation of iron levels in the cell and in ferroptosis. On the other hand, previous work revealed a putative metal binding site in the structure of MEMO1 (15) and demonstrated that it has copper-dependent redox activity (7). Copper binding to MEMO1 has been demonstrated recently (14). Taken together, these findings prompted us to investigate metal binding properties of MEMO1. We expressed MEMO1 as a fusion with the chitin binding domain and intein and purified the protein by chitin affinity chromatography combined with intein self-cleavage, followed by size exclusion chromatography, producing highly pure, well folded protein, as evidenced by NMR spectroscopy (Fig. S11).

Next, we investigated metal binding to pure MEMO1, in the presence of a mixture of the reduced (GSH) and oxidized (GSSG) glutathione (9.5 mM GSH and 0.5 mM GSSG), approximating redox environment in the cytosol (35). Inductively coupled plasma mass-spectrometry (ICP-MS) showed that, under these conditions, MEMO1 binds stoichiometric amounts of iron (approximately 1:1 molar ratio). In the absence of iron, we have also observed binding of copper (added as Cu(I) to MEMO1 at a molar ratio of copper to protein of approximately 0.4. Under oxidative conditions, without glutathione, or other reducing agents, copper, added as Cu (II), bound to MEMO1 at approximately 0.7 molar ratio.

To determine the metal binding affinity, we measured dissociation constants for iron and copper by isothermal titration calorimetry (ITC). The measured *K_d_* value for iron in the presence of glutathione was 5.0 ± 2.6 × 10^-6^ M (Fig. 5A) consistent with the value of 2.4 ± 1.0 × 10^-6^ M determined by microscale thermophoresis (Fig. S12). Glutathione itself binds iron with a *K_d_* of about 10^-5^ M (36), and therefore MEMO1 affinity for free Fe^2+^ ions may be much higher. However, the apparent *K_d_* value measured in the presence of an abundant intracellular low-affinity iron acceptor, such as glutathione, is physiologically more relevant. Copper (I) binding was also detected by ITC, with an estimated *K_d_* of about 3 × 10^-6^ M (Fig. S13).

**Fig. 5.**
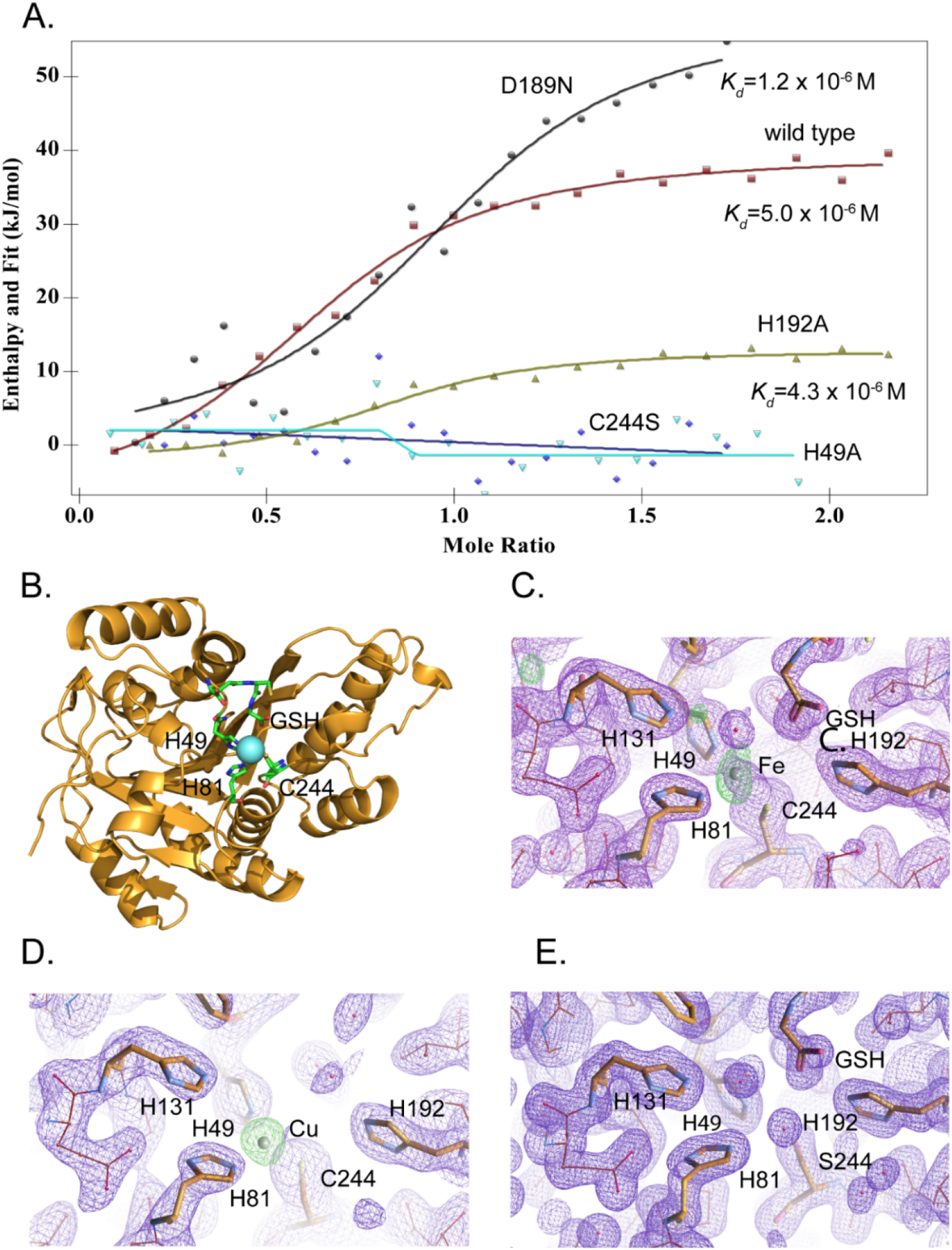
MEMO1 binds iron or copper in the site formed by H49, H81 and C244. **(A)** Iron binding to the wild type MEMO1 and metal binding site mutants analyzed by isothermal titration calorimetry. Dissociation constant (*K_d_*) values are shown under the fitted binding curves for the wild type (*squares*), D189N (*circles*), and H192A (*triangles*) MEMO1 variants. The C244S (*diamonds*) and H49A (*inverted triangles*) variants did not bind iron. **(B)** Structure of the wild type MEMO1 with iron (PDB ID 7KQ8). **(C, D)** Anomalous difference electron density maps showing iron (7KQ8) **(C)** and copper (7L5C) **(D)** coordinated by H49, H81 and C244. The H131 and H192 residues, albeit close to the metal binding site, do not participate in metal coordination. GSH is glutathione. **(E)** Region of the electron density map of the C244S-MEMO1 (7M8H) corresponding to the metal binding site in the wild type (cf. panels C and D).

To reveal the binding mode of each metal, we have crystallized MEMO1 in the presence of iron or copper under the conditions similar to those used previously for solving the structure of the metal free protein (15). To prevent metal oxidation, MEMO1 crystallization with iron (II) or copper (I) was set up in an anaerobic chamber and the plates were incubated under argon, while the crystals were growing. We have solved the structures of MEMO1 with iron (Fig. 5B) at the resolution of 2.15 Å and copper at 2.5 Å, by molecular replacement using the previously solved structure of the metal free protein (PDB ID 3BCZ) (15) (Table S4). The protein fold of both metal-bound forms of MEMO1 is essentially identical to the metal-free structure, with RMSD for α-carbons of the superimposed structures being less than 0.17 Å. In the MEMO1-Fe structure, iron density is clearly visible in the region of the previously predicted metal-binding site with an overall occupancy of 40-60%, as confirmed by the anomalous diffraction map. Iron is coordinated by H49, H81 and C244 (Fig. 5C). As noted previously, these residues correspond to the iron-coordinating residues H12, H61 and E242 in the structure alignment of MEMO1 with catechol dioxygenase LigB (15), to H12, H59 and E239 in the more recent structure of the bacterial gallate dioxygenase DesB (PDB ID: 3WR8), and to H13, H62 and E251 in the aminophenol dioxygenase from *Comamonas sp*. (PDB ID: 3VSH). Iron coordination by two histidines and an aspartate or glutamate is also observed in many other non-heme dioxygenases (37). Thus, iron coordination by two histidines and a cysteine in MEMO1 is an unusual variation on a common theme. A glutathione molecule was also found in the MEMO1-Fe structure, close to the iron binding site, with the glutathione glycine carboxyl forming an electrostatic interaction with H192. This finding is consistent with Fe-GSH binding to MEMO1 observed by ITC. When MEMO1 was crystallized in the presence of copper, the copper atom bound at the same site, and was coordinated by the same residues as iron (Fig. 5D).

To confirm the iron binding site, we generated H49A and C244S mutants, along with H192A and D189N variants, because the latter residues are located in the proximity of the bound iron and were previously proposed to belong to the metal binding site of the protein (7). All the mutant proteins were properly folded as shown by NMR (Fig. S11). As shown by ITC (Fig. 5A), iron binding affinity in the H192A and D189N mutants is not significantly changed, whereas H49A and C244S variants do not bind iron at all. Consistent with the crucial role of C244 in metal binding, no metal density was found in the C244S variant of MEMO1 crystallized in the presence of copper (Fig 5E). Thus, metal coordination in MEMO1 is achieved by H49, H81 and C244, while H192 and D189 do not participate in metal binding in MEMO1.

## Discussion

In summary, our results firmly link MEMO1 to a complex network of iron-dependent processes in the cell. We have shown that MEMO1 is an iron binding protein that exhibits GIs with many other iron-related proteins and regulates iron levels in the cell. Since MEMO1 can bind both iron and copper *in vitro*, the question arises, which metal is bound to MEMO1 in the cell. Our *K_d_* measurements suggest that considerations based on the equilibrium dissociation constants should apply to iron binding in the living cell. Concentration of iron in the labile, i.e. readily exchangeable pool, in the cell is within the low micromolar range (38). Therefore, in the absence of competition with another metal, MEMO1 would bind iron from this pool, as we have demonstrated above for glutathione.

The situation is very different for copper. There is essentially no free or readily exchangeable copper in the cell, as all available copper is tightly bound to proteins with *K_d_* on the order of 10^-13^ M, or less, such as copper chaperones ATOX1 and CCS, metal binding domains of copper ATPases ATP7A and ATP7B, and others (39). Copper transfer between proteins in the cell requires specific protein-protein interactions and is kinetically limited by the rate of such interactions. Therefore, in the absence of specific copper loading mechanisms for MEMO1, it is likely to predominantly bind iron in the cytosol of the living cells, even though copper binding can be observed in *vitro.* Still, consistent with the previous reports (7, 14), MEMO1 may bind copper under oxidative conditions in a specific local environment within the cell, and it is tempting to speculate that metal binding change may trigger a switch between different MEMO1 activities in the cell.

The iron-binding site of MEMO1 is structurally very similar to that of iron-containing extradiol dioxygenases, with a notable difference of a cysteine residue (C244) located at the position occupied by a glutamate in those proteins. No dioxygenase activity has been reported for MEMO1, and we have so far been unable to detect any with a variety of standard substrates we have tried, such as gallate, protocatechuate, 3,4-dihydroxyphenylalanine (L-DOPA), and others. Still, the remarkable structural similarity between MEMO1 and dioxygenases strongly suggests that MEMO1 may catalyze redox reactions involved in biosynthesis or breakdown of a signaling molecule in cancer cells.

We have validated GIs between *MEMO1* and many genes encoding iron-related proteins by studying the effects of loss of function of those gene products on the proliferation of breast cancer and melanoma cell lines with different expression levels of MEMO1. Perhaps the most interesting connection that emerged from these experiments is between MEMO1 and iron transport proteins transferrin and transferrin receptor 2 (TFR2). Both transferrin receptors in human cells, TFR1 and TFR2, are involved in iron transport into the cells. However, whereas TFR1 is an essential protein that accounts for bulk iron uptake, TFR2 plays a regulatory role and appears to protect cells from iron overload (31). Selective activation of high-MEMO breast cancer cell proliferation by TFR2 knockdown, taken together with the high-affinity interactions between MEMO1 and transferrin, and a marked decrease in cytosolic iron concentration in the context of MEMO1 knockout suggest an important role for MEMO1 in maintaining iron homeostasis in cancer cells in conjunction with TFR2 (Fig. 6). MEMO1-dependent activation of cell proliferation by TFR2 knockdown, and the link between MEMO1 and TFR2 expression levels is also observed in melanoma cells, suggesting that MEMO1-TFR2 interaction has a salient role in regulating iron in various cell types.

**Fig. 6.**
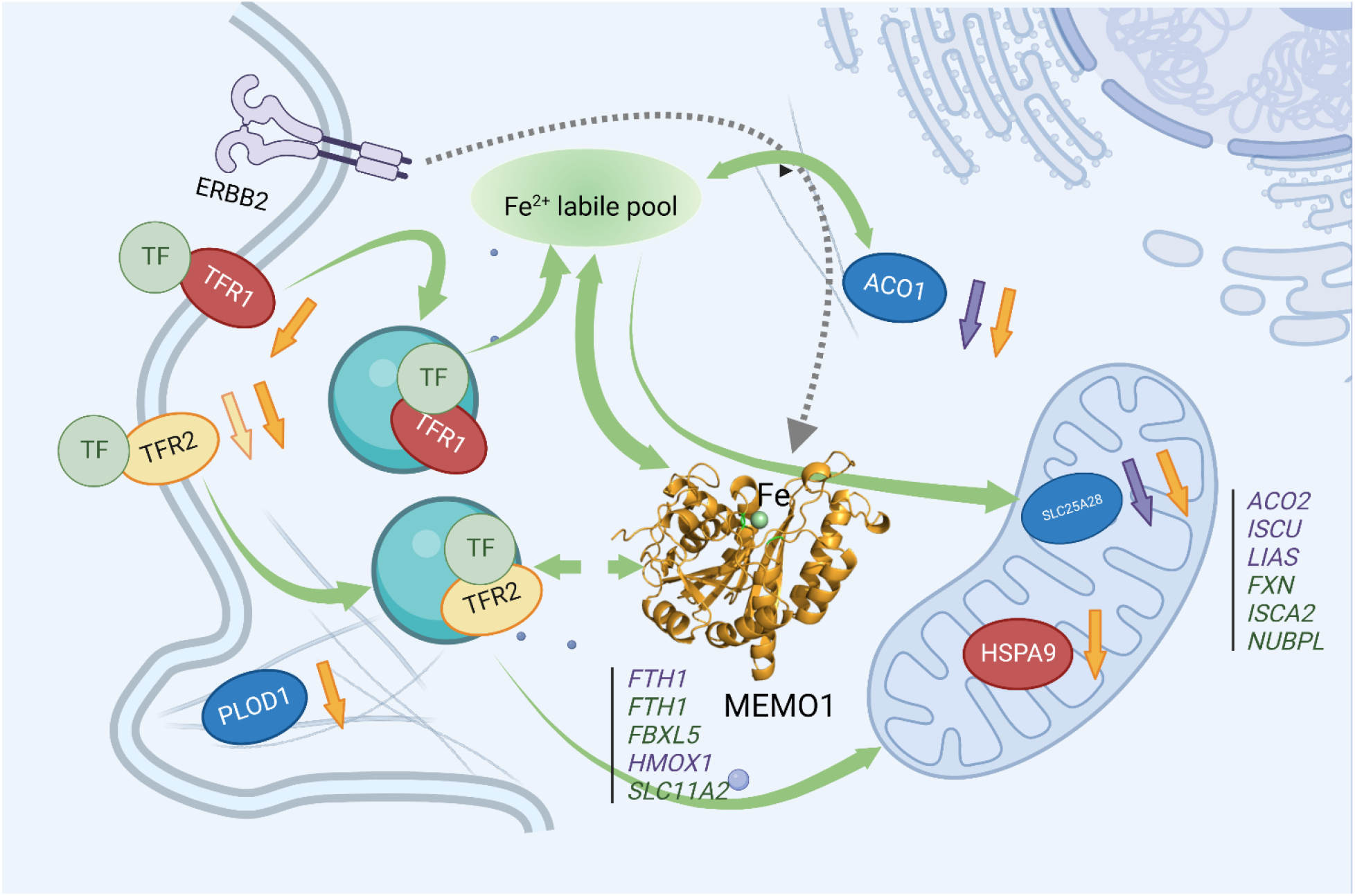
MEMO1 interactions with the other iron-related proteins in breast cancer cells. Proteins involved in the experimentally validated genetic interactions with MEMO1 in the present work are shown as ovals. Other iron-related genes showing interactions with *MEMO1* by database screening are listed in columns, separately for cytosolic and mitochondrial proteins. Proteins or genes shown in *blue* display SDL interactions with MEMO1; those shown in *green* display SL interactions with MEMO1. In *red* are essential proteins; their knockdown severely inhibits cell proliferation regardless of MEMO1 expression status. TFR2 (*yellow*) knockdown uniquely stimulates proliferation of high-MEMO1 cells. Curved *green* unidirectional arrows show iron transport pathways. Curved *green* bidirectional arrows show iron-dependent regulation. Dashed *grey* arrow indicates MEMO1 interaction with ERBB2 receptor. Short up or down arrows indicate reciprocal effects of the MEMO1 and the interacting genes knockouts and knockdowns: protein expression change in low/no MEMO1 cells is indicated by the same color as the protein, MEMO1 expression change with the protein knockdown is indicated by the *orange* arrow. TF-transferrin.

The link between MEMO1 and iron homeostasis is further supported by the GOF-GI between MEMO1 and the iron-dependent regulatory protein ACO1. ACO1, in its iron-free state, functions as an iron-response protein (IRP1): it binds to the iron-responsive elements (IREs) in the 3’-untranslated regions of mRNA encoding iron transporters TFR1 and DMT1, and in the 5’-untranslated regions of mRNA encoding several other iron dependent proteins, including mitochondrial aconitase ACO2, iron efflux protein FPN1 and iron storage protein ferritin (FTN and FTH subunits) (40). Upon iron binding, IRP1 dissociates from mRNA and becomes a functional cytosolic aconitate hydratase (aconitase). IRP1 dissociation results in the destabilization of TFR1 mRNA and a decrease in iron uptake (41).

The selective suppression of the high-MEMO1 cells proliferation by ACO1 knockdown indicates dysregulation of MEMO1-dependent aspects of iron homeostasis in the cell. One of the possible explanations or this finding is a need for a tighter control of iron homeostasis in high-MEMO1 and high-iron cells compared with the cells expressing less MEMO1 and containing less iron. The link between MEMO1 and TFR2, transferrin, and ACO1 would suggest a possible interaction between MEMO1 and TFR1. However, TFR1 knockdown strongly inhibited proliferation of all cell lines, regardless of MEMO1 expression levels, consistent with the essential role of TFR1 in iron uptake. Several other genes regulated by ACO1, including DMT1, FTH1, and ACO2 were detected in our database screening, indicating that MEMO1 is an integral part of iron regulatory network of genetic interactions in the cell.

The MEMO1 iron connection trail leads to mitochondria, as the major intracellular iron consuming organelle. Iron is a cofactor of the highly abundant electron transport proteins in the respiratory chain and several tricarboxylic acid cycle enzymes. It is, therefore, logical to expect that a disruption of iron homeostasis caused by MEMO1 knockout will affect mitochondrial functions. Indeed, we observed that an iron chelator, deferoxamine, causes major changes in the mitochondrial morphology of MEMO1 knockout cells, at a concentration that does not affect the high-MEMO1 parental cells. Perinuclear mitochondrial clustering is one of manifestations of hypoxia (42), which, in turn, may be caused by iron deficiency (40, 43). Absence of the perinuclear clustering in the mitochondria of high-MEMO1 cells combined with higher iron level compared to MEMO1 knockouts suggests that MEMO1 specifically facilitates iron transport into mitochondria. MEMO1 overexpression may help to maintain normal metabolism of cancer cells by increasing iron levels in mitochondrial under hypoxic conditions that are frequently found in tumors.

Remarkably, transferrin and TFR2, which showed physical or genetic interactions with MEMO1 in our work, have been previously reported to be a part of the novel iron transport pathways to mitochondria in substantia nigra dopamine neurons (44) and in erythroid progenitor cells (45). Although it remains to be determined, whether a similar pathway exists in breast cancer cells that we studied, our data provide the first indication that MEMO1, TFR2 and transferrin may all be components of the same iron transport pathway to mitochondria.

Several proteins involved in the biosynthesis of iron-containing cofactors in mitochondria showed genetic interactions with MEMO1 in our genome-wide *in silico* screening. SLC25A28 (Mitoferrin-2), which mediates iron uptake in mitochondria, displayed GOF-GI with MEMO1, its knockdown selectively suppressing proliferation of MEMO1-overexpressing breast cancer cells. This may indicate that dysregulation of iron transport processes in high-MEMO1 cells makes them more susceptible to ferroptosis by diverting iron off pathway, to participate in the damaging oxidative reactions. In fact, in our experiments, high-MEMO1 breast cancer cells were significantly more sensitive to the ferroptosis inducer RSL3 than MEMO1 knockouts.

Genetic interactions between MEMO1 and PLOD1 in breast cancer cells that we found are particularly notable. PLOD1 is highly expressed in many malignant tumors, likely contributing to the epithelial to mesenchymal cell transition in the course of cancer progression (46). The MEMO1-PLOD1 genetic interaction establishes a direct connection between iron, MEMO1 and cancer cell motility. PLOD1 is an iron-containing enzyme, which participates in the collagen assembly and in the regulation of collagen synthesis and extracellular matrix remodeling, consistent with the established role of MEMO1 in cancer cell tissue invasion and metastasis.

Many of the GIs between MEMO1 and the other iron-related proteins that we have investigated in the present work manifest themselves not only in MEMO1-dependent effects of the second gene knockdown on cell proliferation, but also in the connections between the expression levels of the two proteins. Such connections are revealed both in the weak but statistically highly significant correlations between the expression levels of MEMO1 and TFR1, TFR2 and PLOD1 across multiple breast cancer cell lines and in the reciprocal effects of gene knockdowns and knockouts on protein expression observed in our experiments. Thus, TFR2 and SLC25A28 (mitoferrin-2) levels are markedly decreased in breast cancer cells with MEMO1 knockdown and knockout. Conversely, TFR1, TFR2, SLC25A28 and PLOD1 knockdowns decrease expression levels of MEMO1. These correlations indicate coregulation of expression of many iron-related genes and *MEMO1* (Fig. 6).

Across the board comparison of MEMO1 genetic interactions and MEMO1 knockout effects between breast cancer and melanoma cells indicates that variations in MEMO1 levels overall have stronger effects in breast cancer cells than in melanoma. Therefore, MEMO1 overexpression relative to the normal tissue rather than absolute MEMO1 levels in the cell appears to be a hallmark of hypersensitivity to iron homeostasis disruption.

In summary, our work has revealed that MEMO1 is an iron-binding protein that regulates iron homeostasis in cancer cells. MEMO1 overexpression may help to maintain normal metabolism of cancer cells by increasing iron levels in mitochondrial under hypoxic conditions. Thus, MEMO1 may serve as a biomarker of tumors particularly sensitive to the therapies targeting iron metabolism in the cell. Genetic interactions of MEMO1 may be targeted to suppress metastasis in breast cancer and other malignancies with high MEMO1 expression level. MEMO1 structure and iron coordination mode suggest that it may be involved in the biosynthesis or processing of a signal molecule in the cell.

## Methods

### Generation of Memo1 knockout and knockdown cell lines

MDA-MB-231 (breast cancer) and A-375 (melanoma) cell lines were obtained from ATCC and cultured in Dulbecco’s Modified Eagle Medium (DMEM, HyClone) supplemented with 10% Fetal Bovine Serum (FBS, Gibco). MEMO1 knockdowns and knockouts were generated using CRISPR/Cas9 TrueGuide synthetic crRNA technology (*Invitrogen*) with the guiding crRNA targeting MEMO1 exons 4-6. Genomic cleavage by Cas9 was confirmed using GeneArt Genomic Cleavage Detection kit (*Life Technologies*) to amplify the region of genomic DNA targeted for cleavage. Individual clones containing MEMO1 knockouts and knockdowns were generated by limiting dilution and tested for MEMO1 expression by Western blot.

### Western blotting

Cells were scraped, washed in PBS, resuspended in lysis buffer (50 mM HEPES, pH 7.4, 150 mM NaCl, 0.2% Triton X-100, cOmplete ULTRA Protease Inhibitor Cocktail (*Roche*) and incubated on ice for 30 min. Nuclei and cell debris were separated by centrifugation at 6,000 g for 10 minutes. Protein concentration in the supernatant was measured by BCA (*Pierce*). Proteins were separated using 4-20% Mini-Protein TGX Precast gels (*Bio-Rad*) and transferred to 0.2 μm nitrocellulose membrane (*Bio-Rad*). Membranes were blocked with 5% Amersham ECL Blocking Reagent (*GE*) in PBS with 0.1% Triton X-100 (PBST) for 1 hour at room temperature, then incubated with the primary antibodies at 4°C overnight (see Supplementary Information for the detailed antibody and treatment description). After washing and incubation with the secondary antibody, membranes were washed with 0.5% blocking reagent in PBST (3 times for 5 minutes) and imaged using standard ECL solutions and G:BOX (*Syngene*).

### Genome wide in-silico screening

Using a previously described concept (25), we have screened the Marcotte et al (47) and Achilles Project databases (24), and Project DRIVE (27) RNAi datasets and CERES (28, 29) CRISPR-Cas9 dataset. The cell lines in each dataset were classified based on the expression of MEMO1 from Cancer Cell Line Encyclopedia (CCLE) (48) database. The difference in the gene essentiality score between the top 5 % of high MEMO1 expressing cell lines and bottom 5 % of low MEMO1 cell lines for pan-cancer and top 25 % of high MEMO1 expressing cell lines and bottom 25 % of low MEMO1 cell lines for breast cancer cell lines were calculated and ranked statistically significant hits (Wilcoxon rank-sum test *P<0.05*) by the difference in median values of the essentiality scores. We thus generated two types of datasets, one containing the genes essential in MEMO1-low cell lines, the other in MEMO1-high cell lines. Collectively, these datasets contained results from 1028 cancer cell lines, including 92 breast cancer cell lines. Enrichment analysis for the SL/SDL interaction partners were performed using Gene Set Enrichment Analysis (GSEA) software (49).

### The shRNA knockdown assays

MEMO1 genetic interactions predicted by genome wide gene knockout and knockdown database analysis were validated by measuring proliferation rates of cells with high-level MEMO1 expression (parental cell lines MDA-MB-231 and A-375), MEMO1 knockdowns (M67-2 and A67-4 respectively) and complete MEMO1 knockouts (M67-9 and A67-16 respectively) with shRNA knockdowns of PLOD1, HSPA9, SLC25A28, TFR1, TFR2 or ACO1, compared to the control RFP-shRNA. Pooled shRNAs targeting each of the tested genes were delivered into the cells by lentiviral transfection. In brief, cells were transfected with lentivirus and incubated for 24 hours, then the virus was removed, and cells were incubated with puromycin for the next 48 hours, then trypsinyzed and plated onto 96-well plates in the replicates of 8 with at the density of 1,000 cells per well for A-375, A67-4 and A67-16 cells, and 2,000 cells per well for MDA-MB-231, M67-2 and M67-9 in the presence of puromycin. Cells were imaged using Incucyte S3 (*Sartorius*) every 8 hours for the next 120-140 hours. Cell confluency was measured and used to calculate proliferation rates by fitting data using Logistic Growth equation with GraphPad Prizm v. 9. The rest of the cells were plated onto 100 mm plates and harvested for Western blot analysis after 48 hours.

### Cytotoxicity assay

Cytotoxicity of RSL3 was measured using resazurin assay. Cells were plated onto 96-well plates at the density of 1,000 cell per well for MDA-MB-231 cell line and its derivatives and 750 cell per well for A-375 cell line and its derivatives. The next day cells were titrated with RSL3 and incubated for the next 48 hours. After incubation, media was discarded and replaced by the fresh one containing 88 µM resazurin (*Sigma-Aldrich*). Cells were incubated overnight, and fluorescence was measured using 540 nm excitation and 590 nm emission.

### Malondialdehyde (MDA) assay

Cells were grown on 60 mm plates to 80% confluency, harvested and resuspended in 1 ml of cold PBS. A 100 µl volume of suspension was separated for measuring protein concentration by BCA (Pierce). Cells were pelleted at 14,000 g for 1 min, and lipid peroxidation was measured using MDA assay kit (*Abcam*) according to manufacturer’s instructions.

### Immunofluorescence microscopy

Cells were plated onto glass coverslips at the confluency ∼40-50% and incubated overnight. The next day 1 µM of DFX was added and cells were incubated overnight. Following the incubation, cells were washed with cold PBS and fixed using 50:50 mix of methanol and acetone (−20°C) for 30 s, blocked in 5% BSA in PBS overnight and incubated with anti-GRP75 antibodies (D-9, *Santa Cruz Biotechnology*) for 1 h at room temperature. Cells were washed three times with PBS and incubated with secondary goat anti-rabbit antibody labeled with Alexa Fluor 633 (*Invitrogen*) in the dark for 1 h at room temperature. After wash in PBS, cells were mounted onto the glass microscope slides using ProLong Diamond antifade mountant with DAPI (*Invitrogen*) and imaged using Leica Dmi8 confocal microscope after 48 hours.

Imaging with MitoTracker CM-H2Ros was performed using live cells: after incubation with DFX the media was removed and substituted with 200 nM MitoTracker solution in DMEM (no FBS), cells were incubated for 30 min, then media was discarded, and cells were incubated for 5 min in DMEM without serum. Following that, cells were washed with PBS and incubated in 8 µM Hoechst 33422 (*Invitrogen*) for 10 min, rinsed with PBS, placed in Live Cells Imaging Solution (*Molecular Probes*) and imaged on Leica Dmi8 confocal microscope within 2 hours post staining.

### ICP-MS measurements

Cells were grown on 100 mm plates (6 biological replicates), rinsed with PBS, trypsinyzed, then resuspended in 1 ml of cold PBS. A 0.1 ml volume of suspension was kept for determination by BCA (Pierce), the rest was pelleted at 14,000 g for 1 min. Cell pellets were resuspended in 0.4 ml of PBS and lysed by 25 passages through a 27-gauge syringe needle. Cell lysates were centrifuged for 2 minutes at 1,000 g to remove remaining whole cells and nuclei, and the supernatant was centrifuged again at 10,300 g for 10 min. Supernatant containing the cytosolic fraction was collected and centrifuged for 30 min at 100,000 g. Pellet containing crude mitochondrial fraction was resuspended in 0.5 ml of PBS and centrifuged again, then resuspended in 50 μl PBS. Protein concentration in the samples was determined by BCA assay (Pierce). Prior to ICP-MS analysis, the cytosolic fraction was diluted with 1% nitric acid (trace metal grade, Thermo Fisher Scientific). The mitochondrial fraction was briefly digested with concentrated nitric acid (trace metal grade, Thermo Fisher Scientific) at 90°C and subsequently diluted with 1% nitric acid.

ICP-MS measurements were performed using an Agilent 7700x equipped with an ASX 500 autosampler in the OHSU Elemental Analysis Shared Resource (Oregon, USA). Data were quantified using weighed, serial dilutions of a multi-element standard (CEM 2, (VHG Labs VHG-SM70B-100) Fe, Cu, Zn) and a single element standard for Ca (inorganic ventures CGCA1) and P (VHG Labs, PPN-500).

### Statistical Analysis

All experiments were repeated at least 3 times; the results are presented as averages ± standard deviation (SD). Statistical significance for the difference between the datasets was calculated using one-way ANOVA with follow-up Tukey’s multiple comparison tests or, when appropriate, two-way ANOVA with follow-up Sidak’s multiple comparison tests. All calculations were carried out using GraphPad Prizm v. 6. The *P*-values below 0.05 were considered statistically significant. Statistical significance level in the figures is indicated as * or # for *P*˂0.05, ** or ## for *P*˂0.01, *** or ### for *P*˂0.001, and **** or #### for *P*˂0.0001.

### Protein expression and purification

DNA sequence encoding MEMO1 was codon optimized for *E. coli* expression and prepared by chemical synthesis (*Integrated DNA Technologies, Inc.*). Mutant variants of MEMO1 were generated by site-directed mutagenesis. MEMO1 was expressed as fusion with the chitin-binding domain and intein using vector pTYB12 (*New England BioLabs*) in *E. coli* BL21(DE3) and purified by chitin affinity chromatography combined with intein self-cleavage, essentially as described previously (50). MEMO1 was additionally purified by size exclusion chromatography on a Superdex 75 10/300 GL Increase column (*GE Life Scienc*es) in a buffer containing 50 mM HEPES-Na, pH 7.4, 150 mM NaCl and 0.6 mM *tris*-(2-carboxyethyl)phosphine and concentrated by membrane filtration.

### Isothermal titration calorimetry

For ITC experiments, MEMO1 was dialyzed against 50 mM HEPES, pH 7.4, 150 mM NaCl, 9.5 mM reduced glutathione (GSH), 0.5 mM oxidized glutathione (GSSG) under argon. Iron (II) sulfate was dissolved in the used dialysis buffer. ITC was performed on a TA Instruments (New Castle, DE) Low Volume Nano calorimeter using ITCRun software and analyzed with NanoAnalyze software (TA Instruments) using independent model setting. A 0.5 mM solution of iron sulfate was titrated into 0.17 ml of 25 μM MEMO1, 1.5-2.5 μl per injection at 300 s intervals at 25°C under constant stirring at 300 r.p.m. Dilution heat correction was applied by titrating iron sulfate into the ITC buffer containing no protein. Most of the titrations were performed in duplicate or triplicate.

### Microscale thermophoresis

Purified MEMO1 was labeled with Red-NHS dye (*NanoTemper Technologies GmbH*) and diluted in 10 mM MES, pH 6.0, 250 mM NaCl, 0.1% Pluronic acid. Human holo- and apo-transferrin (*Sigma-Aldrich*) were dissolved the same buffer at a concentration of 0.1-0.15 mM and titrated into 20 nM labeled MEMO1. Measurements were performed on a NanoTemper Monolith NT.115.

### Protein crystallization

Protein crystals were obtained by mixing 0.5 µl of purified MEMO1 (12-13 mg/mL) with 0.5 µl mother liquor containing 100 mM 2-(N-morpholino) ethanesulfonic acid (MES) pH 5.5-7.0 and 22.5 % PEG-3350 and incubated at 20 °C. Protein crystals that contained reduced Fe^2+^ and Cu^+^ were set up inside an anaerobic chamber with 3 ppm of O_2_ in the atmosphere of 4 % H_2_/N_2_ mixture. Mother liquor containing 1 mM metal was incubated inside the chamber with mixing to get rid of oxygen dissolved in solution prior to crystallization. Crystals were harvested and stored in liquid nitrogen using 20 % glycerol as cryoprotectant. Crystals diffracted to 1.75 Å – 2.55 Å and belonged to orthorhombic system, space group P2_1_2_1_2, with cell dimensions of approximately *a*=140 Å, *b*= 87 Å, *c*=98 Å, α=β=γ=90°, and contained four molecules in the asymmetric unit.

### Data collection and structure refinement

Diffraction data were collected at the Canadian Light Source, CMCF section, using beamlines 08ID and 08BM and Pilatus 6M detector. Data were integrated and scaled using XDS package to 2.15 Å (for MEMO1-Fe complex), 2.55 Å (for MEMO1-Cu complex) and 1.75 Å (for MEMO1 C244S mutant) (51). Initial phases were obtained using Phaser MR (52) within ccp4i package and previously solved MEMO1 structure (PDB code: 3BCZ) as a model for molecular replacement (15). The final models of MEMO1 were refined using phenix.refine (53) with manual rebuilding using COOT (54). The coordinates and structure factors have been deposited to the Protein Data Bank with ID codes: 7KQ8 (MEMO1-Fe), 7L5C (MEMO1-Cu), and 7M8H (MEMO1 C244S).

### Materials availability statement

MEMO1 knockout and knockdown cell lines and MEMO1 expression vectors are available from the authors upon a reasonable request.

## Acknowledgments

We thank the Canadian Light Source, CMCF section staff for support during data collection. We gratefully acknowledge the use of instrumentation at the Protein Characterization, Crystallization Facility (PCCF) and Phenogenomic Imaging Centre of Saskatchewan (PICS) supported by College of Medicine, University of Saskatchewan. ICP-MS measurements were performed by Marvel Davis in the OHSU Elemental Analysis Shared Resource with partial support from NIH core grant S10RR025512. NMR spectra were recorded at the National Magnetic Resonance at Madison (NMRFAM), which is supported by NIH grant P41 GM103399 and by the University of Wisconsin-Madison. We are grateful to Anjuman Ara and Paria Nouri for assistance with some of the experiments.

## Materials and Methods supplementary information

### Antibody and Western blotting details

The following antibodies were used: anti-Memo1 antibodies (mouse monoclonal antibodies AT1E9, sc-517412, Santa Cruz Biotechnology, dilution 1:500 in 0.5% of blocking reagent on PBST), anti-PLOD1 (LLH1) antibodies (mouse monoclonal antibodies B-5, sc-271640, Santa Cruz Biotechnology, dilution 1:500 in 0.5% of blocking reagent on PBST), anti-OGDH antibodies (rabbit polyclonal antibodies PA5-28195, Invitrogen, dilution 1:2,000 in 0.5% of blocking reagent on PBST), anti-TfR1 antibodies (mouse monoclonal anti-CD71 antibodies 3B8 2A1, Santa Cruz Biotechnology, dilution 1:500 in 0.5% of blocking reagent on PBST), anti-TfR2 antibodies (mouse monoclonal antibodies B-6, sc-376278, Santa Cruz Biotechnology, dilution 1:100 in 0.5% of blocking reagent on PBST), anti-Aco1 antibodies (rabbit polyclonal antibodies PA5-41753, Invitrogen, dilution 1:1,000 in 0.5% of blocking reagent on PBST), anti-SCL25A28 (rabbit polyclonal antibodies against mitoferrin 2, BS-7157R, Bioss, ThermoFisher Scientific, dilution 1:500 in 0.5% of blocking reagent on PBST), anti-Grp75 antibodies (mouse monoclonal anti-HSPA9 antibodies D-9, sc-133137, Santa Cruz Biotechnology, dilution 1:1,000 in 0.5% of blocking reagent on PBST), anti-actin β antibodies (MA5-15452, Invitrogen, dilution 1:10,000 in 0.5% of blocking reagent on PBST). Following incubation with primary antibodies, membranes were washed in 0.5% blocking reagent on PBST three times for 5 minutes, and incubated with the following secondary antibodies for 1 hour at room temperature: Amersham™ ECL™ Anti-Mouse IgG, Horseradish Peroxidase (HRP)-Linked Species-Specific Whole Antibody (from sheep) (NA931V), dilution 1:1,000, or Pierce donkey anti-rabbit HRP-linked antibodies (#PI31458), dilution 1:2,500. Membranes probed with anti-TfR2 antibodies were incubated with m-IgGk BP-HRP secondary antibodies (Santa Cruz, sc-525409, dilution 1:1,000).

### Site-directed mutagenesis details

To generate MEMO1 binding site variants, QuickChange method (Agilent) was used with the following primers:

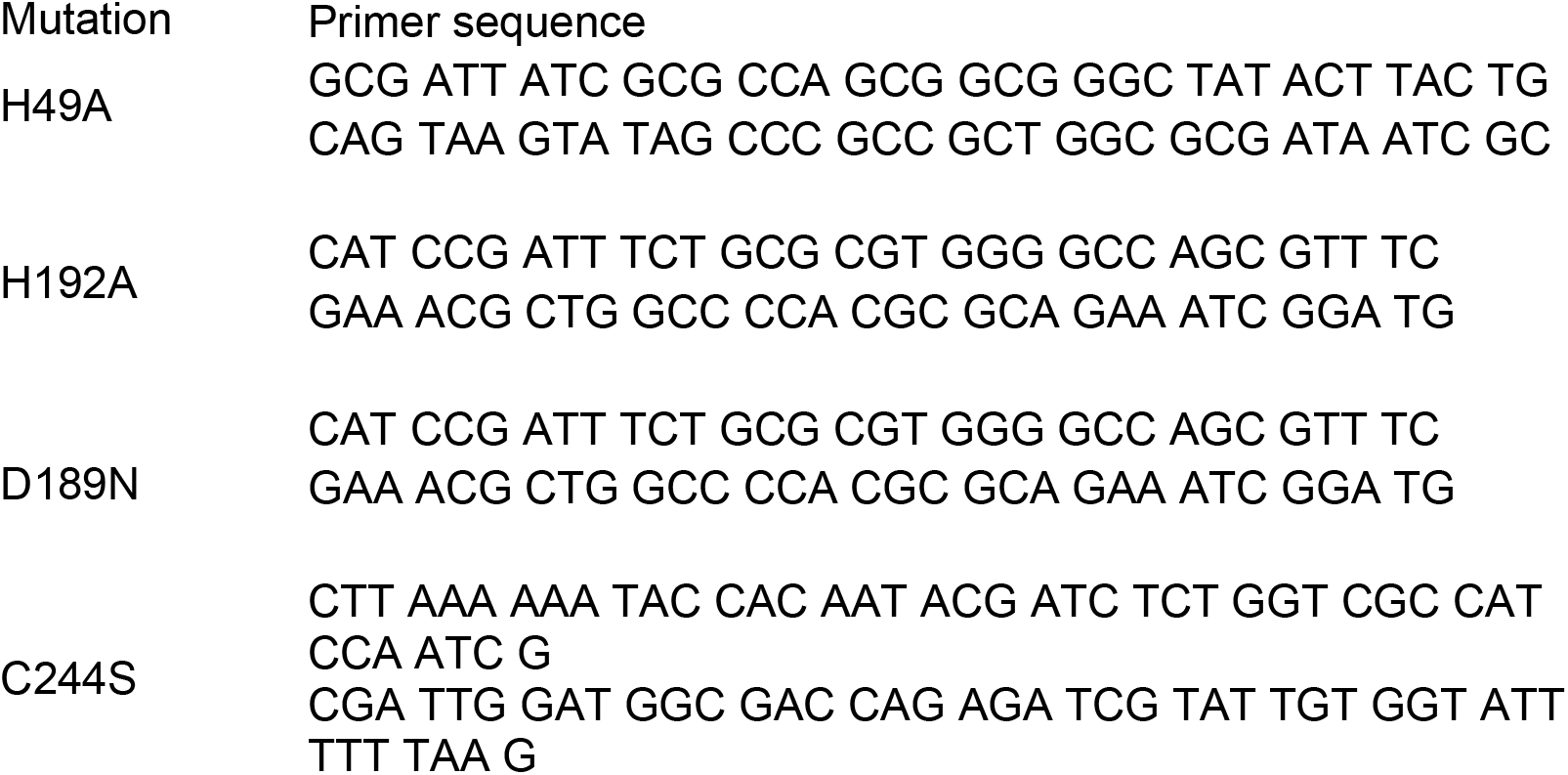

### CRISPR knockout details

To generate MEMO1 knockouts, CRISPR1006367_SGM sgRNA (Thermo Fisher Scientific) was used with the target DNA sequence TACGGAGAACTGTGGAAGAC, and the protospacer adjacent motif AGG. Primers ACATACCCACATACACTCAC and CCTCCTTCCTTTCCTTTCTTTC were used to amplify the region of genomic DNA targeted for cleavage by this sgRNA, and the subsequent genomic cleavage detection.

---

**Fig. S1.**
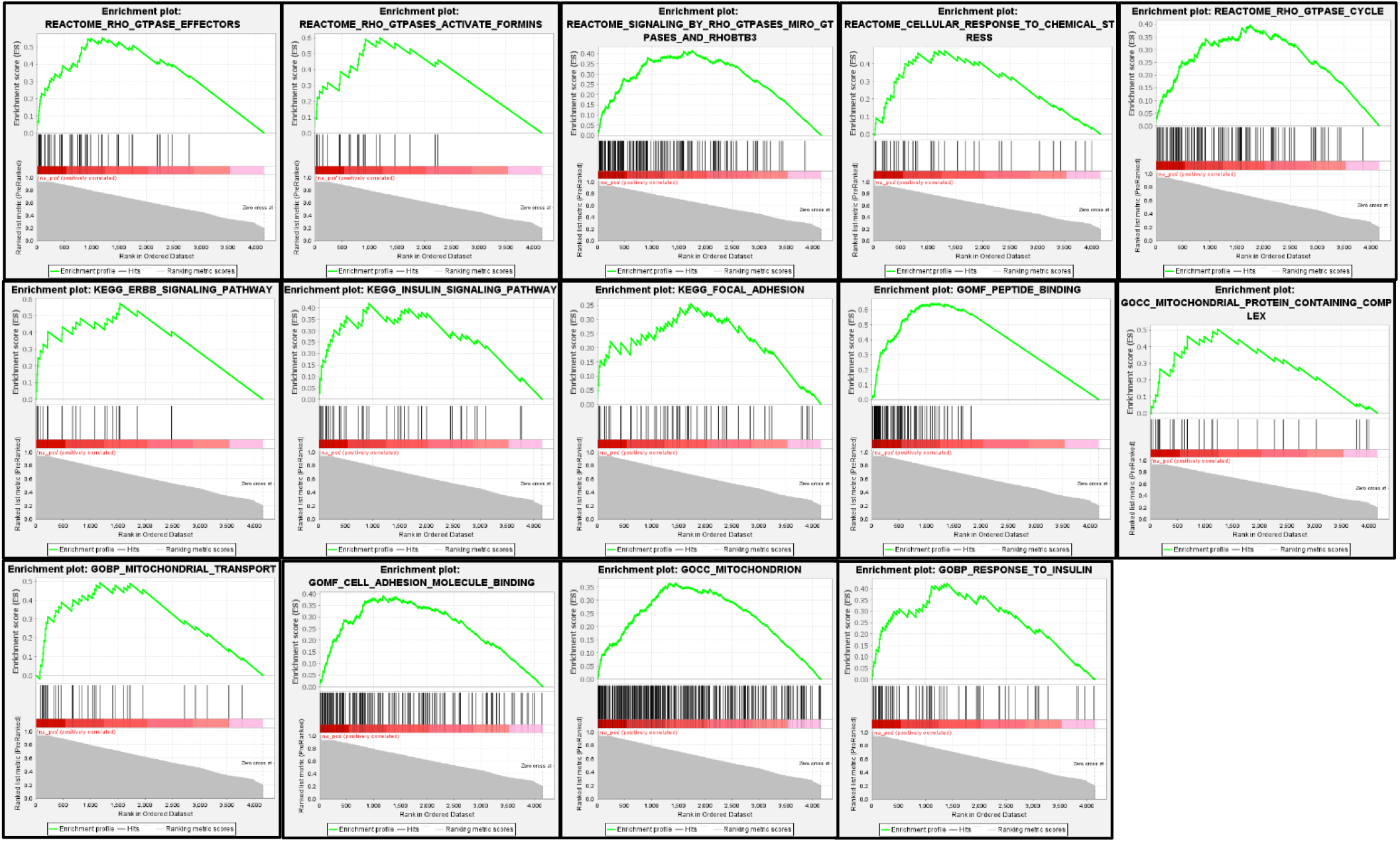
Enrichment plots for MEMO1 loss of function genetic interaction hits.

**Fig. S2.**
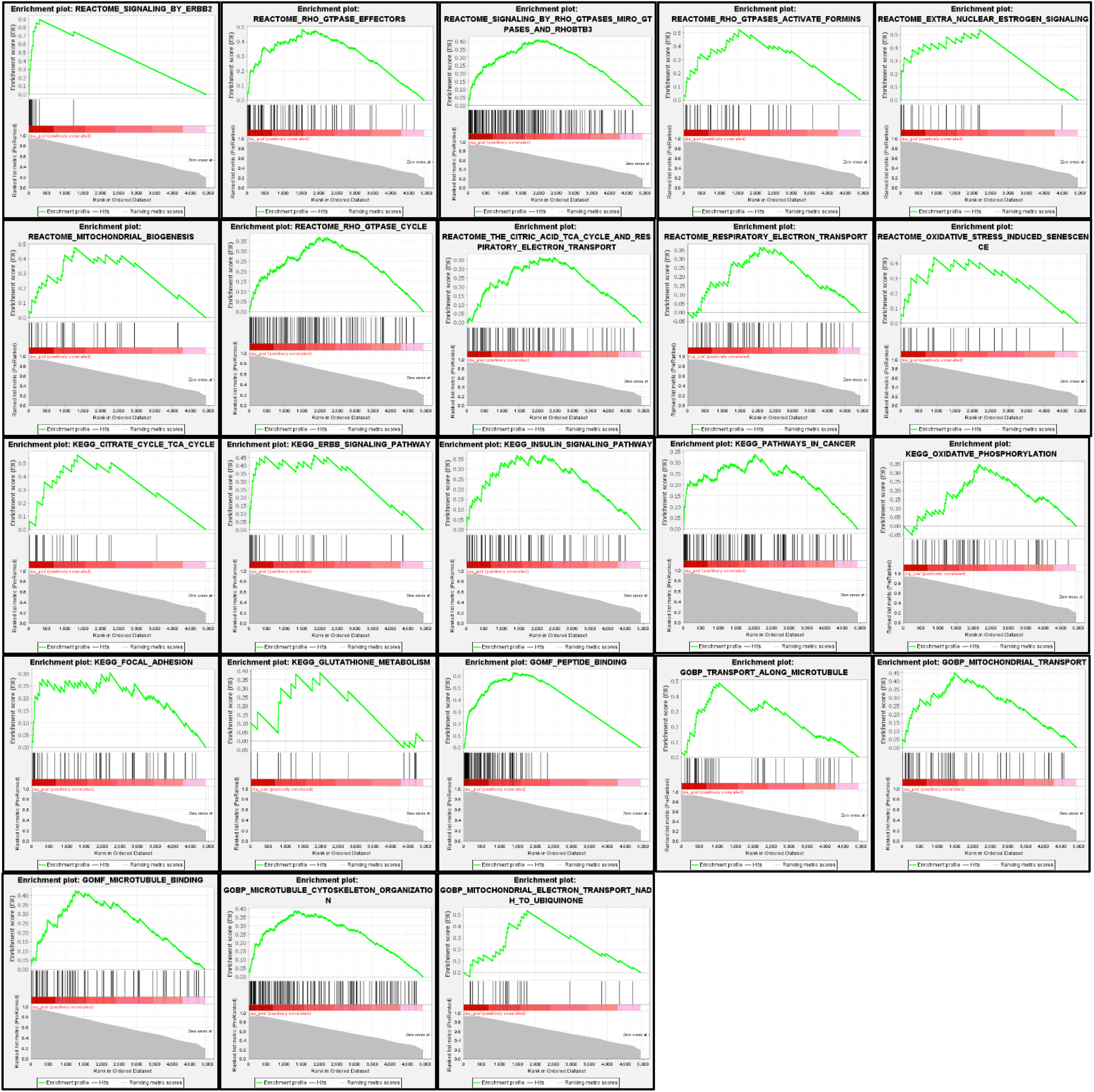
Enrichment plots for MEMO1 gain of function genetic interactions hits.

**Fig. S3.**
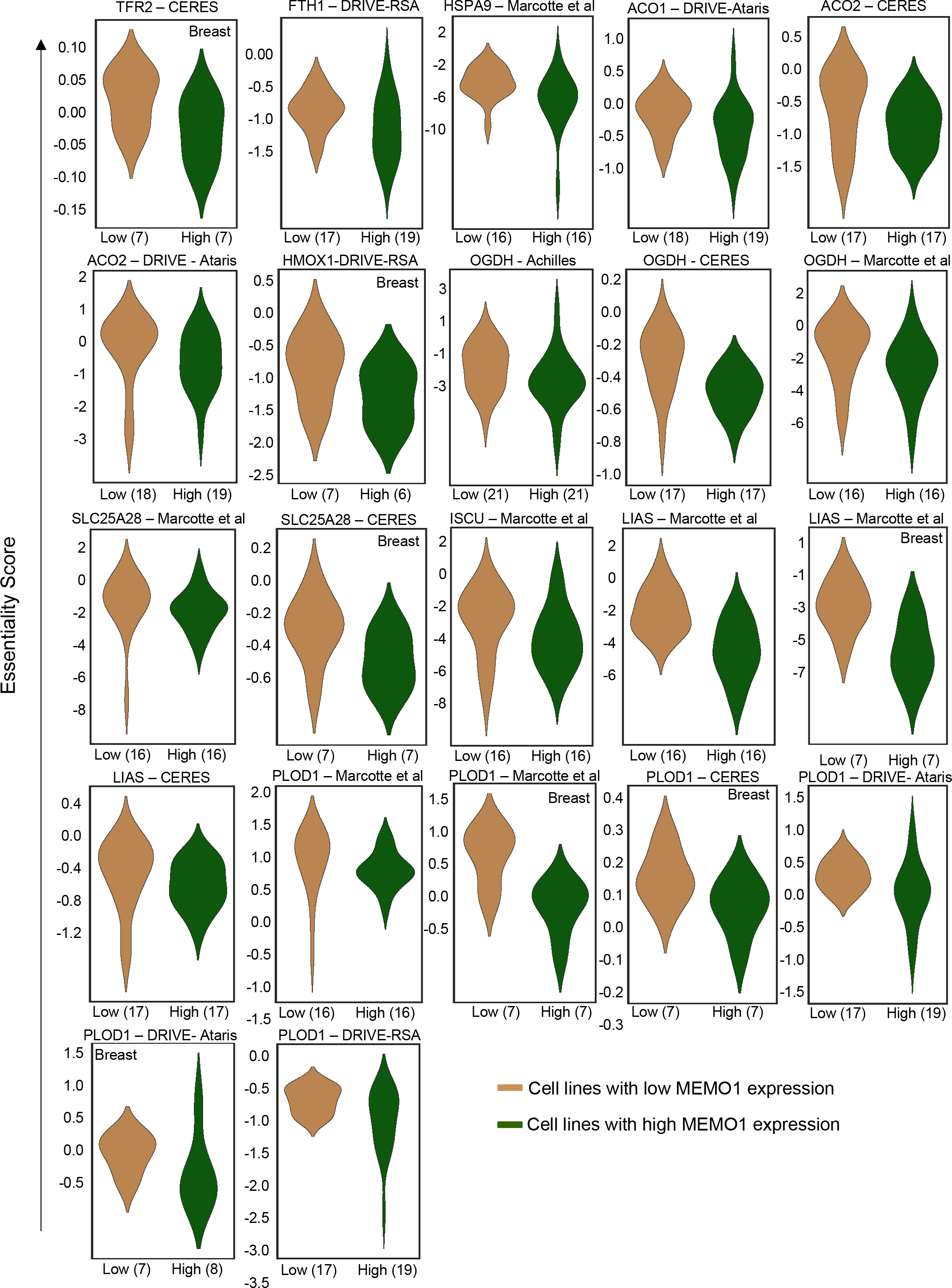
Iron related genes exhibiting GOF interactions with MEMO1. Gene essentiality score distribution in the low- and high-MEMO1 expressing groups with the number of cell lines in each group is shown (cf. Table S1).

**Fig. S4.**
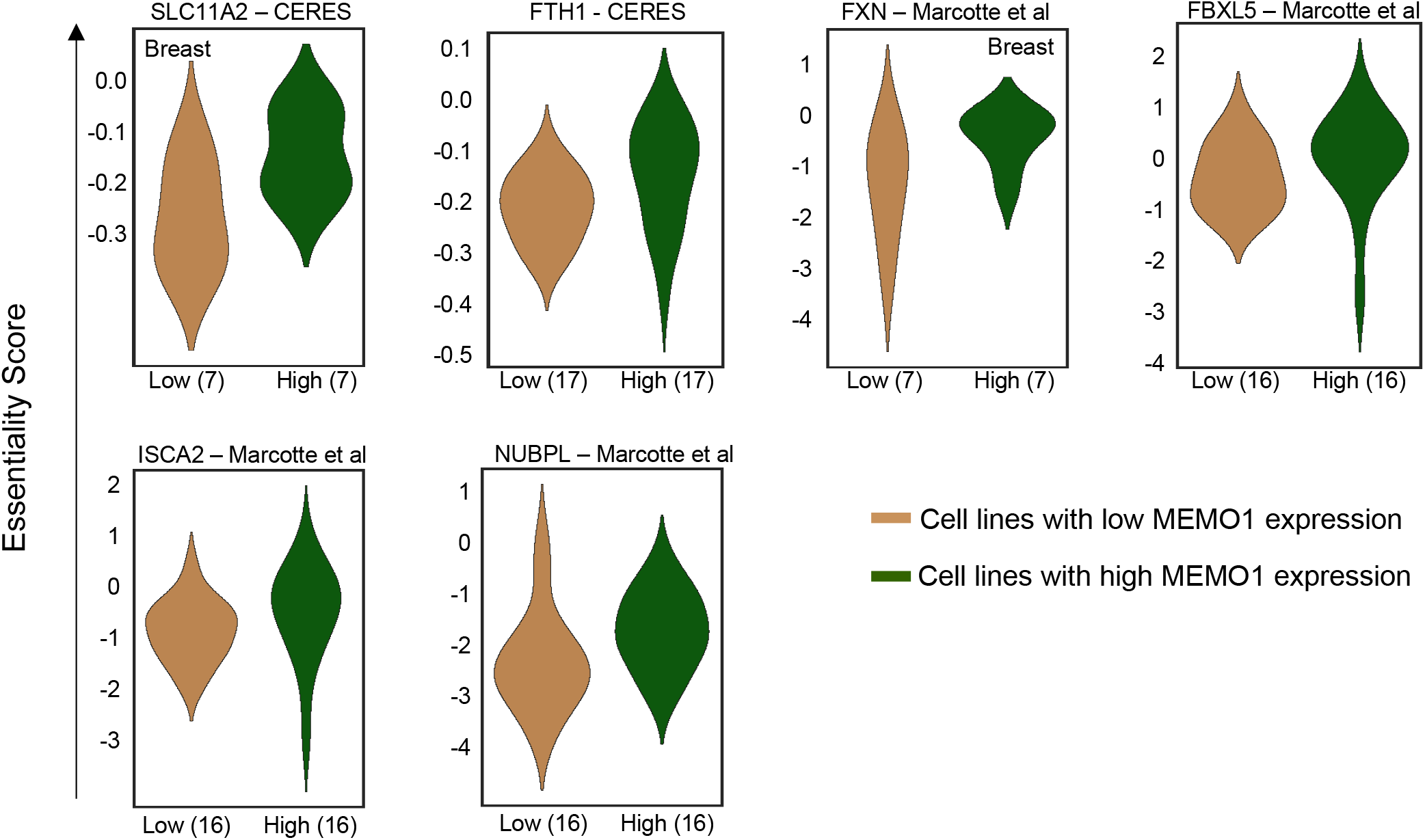
Iron related genes exhibiting LOF interactions with MEMO1. Gene essentiality score distribution in the low- and high-MEMO1 expressing groups with the number of cell lines in each group is shown (cf. Table S2).

**Fig. S5.**
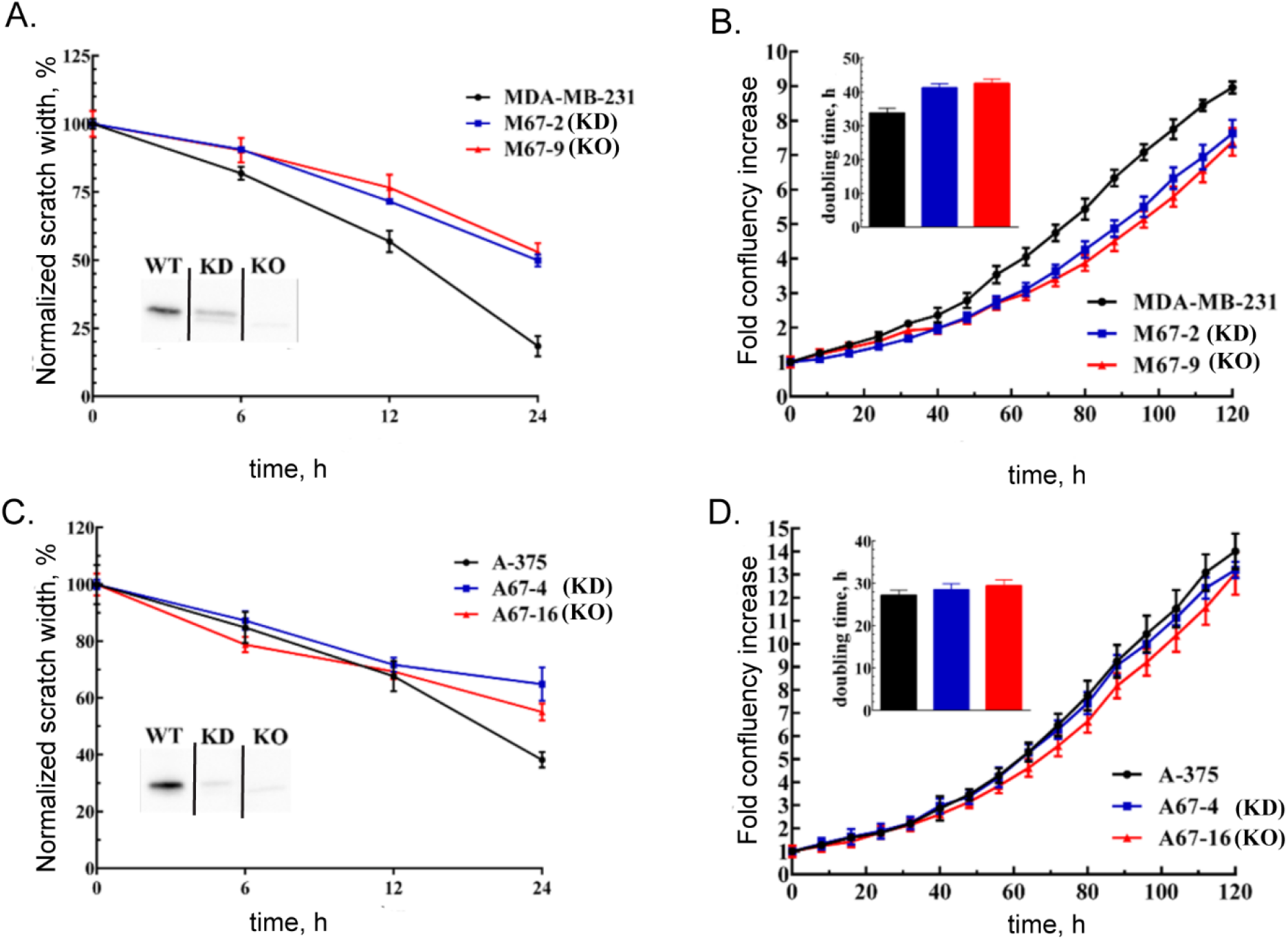
MEMO1 knockout and knockdown in breast cancer and melanoma cell lines result in decrease in cellular motility and cell growth rate. (A) Wound healing assay for breast cancer cell lines MDA-MB-231 (parental), M67-2 (MEMO1 knockdown) and M67-9 (MEMO1 knockout). MEMO1 Western blots demonstrating MEMO1 expression levels in the parental cell line (WT) and in the CRISPR/Cas9 knockdown (KD) and knockout (KO) are shown in the inset. (B) Growth curves of MDA-MB-231, M67-2 and M67-9 cells. (C) Wound healing assay for A-375, A67-4 (MEMO1 knockdown) and A67-16 (MEMO1-knockout) cells. MEMO1 Western blots demonstrating MEMO1 expression levels in the parental cell line (WT) and in the CRISPR/Cas9 knockdown (KD) and knockout (KO) are shown in the inset. (D) Growth curves of A-375, A67-4 and A-67-16 cells.

**Figure S6.**
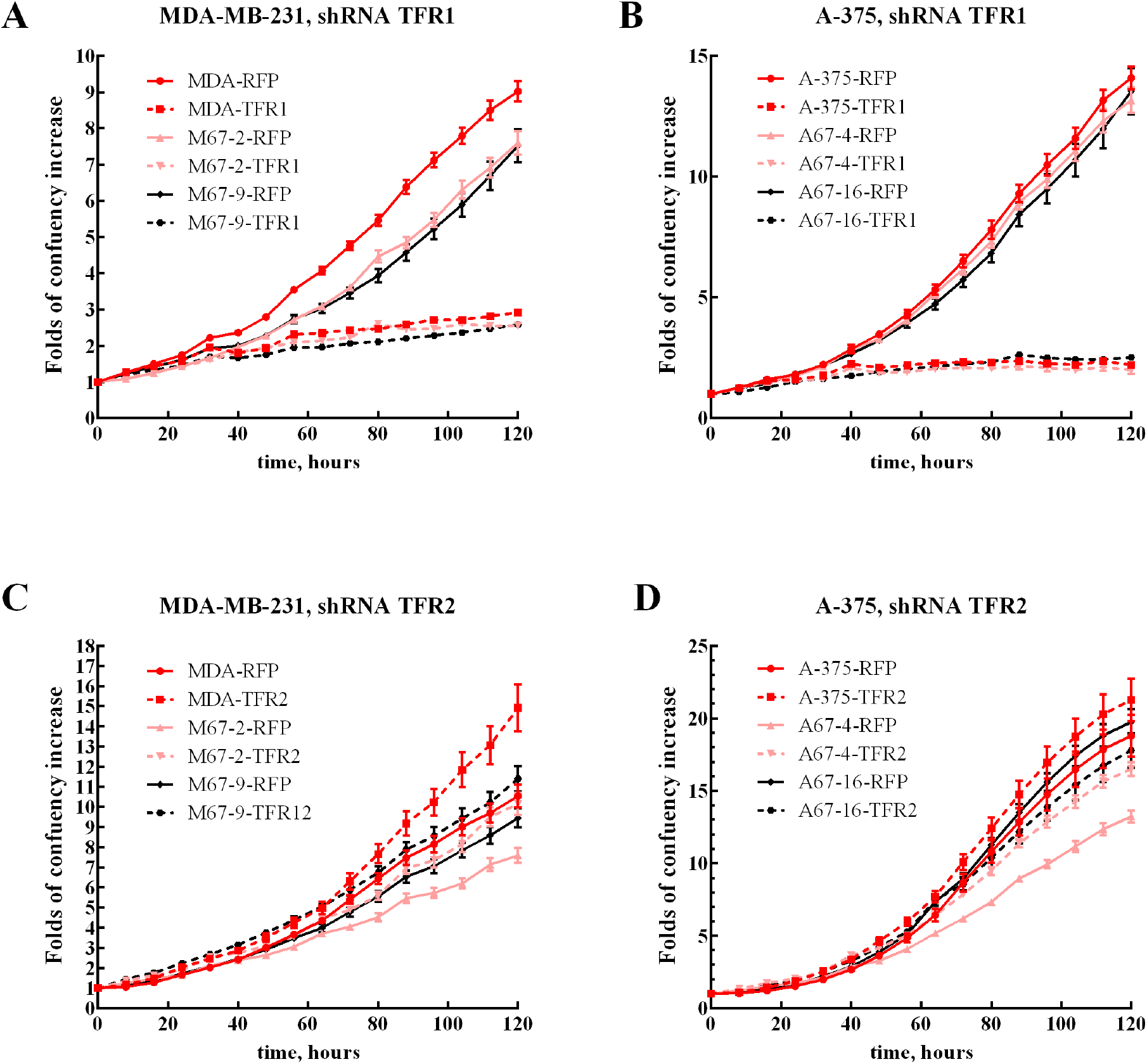
Growth curves of MDA-MB-231 (A, C) and A-375 (B, D) with high expression of MEMO1 (MDA-RFP and A-375-RFP, red solid lines), low expression of MEMO1 (M67-2-RFP and A67-4-RFP, pink solid lines) and no MEMO1 expression (M67-9-RFP and A67-16-RFP black solid lines), in comparison with the same cell lines transduced with TFR1 (A, B) or TFR2 (C, D) shRNA (corresponding dashed lines).

**Figure S7.**
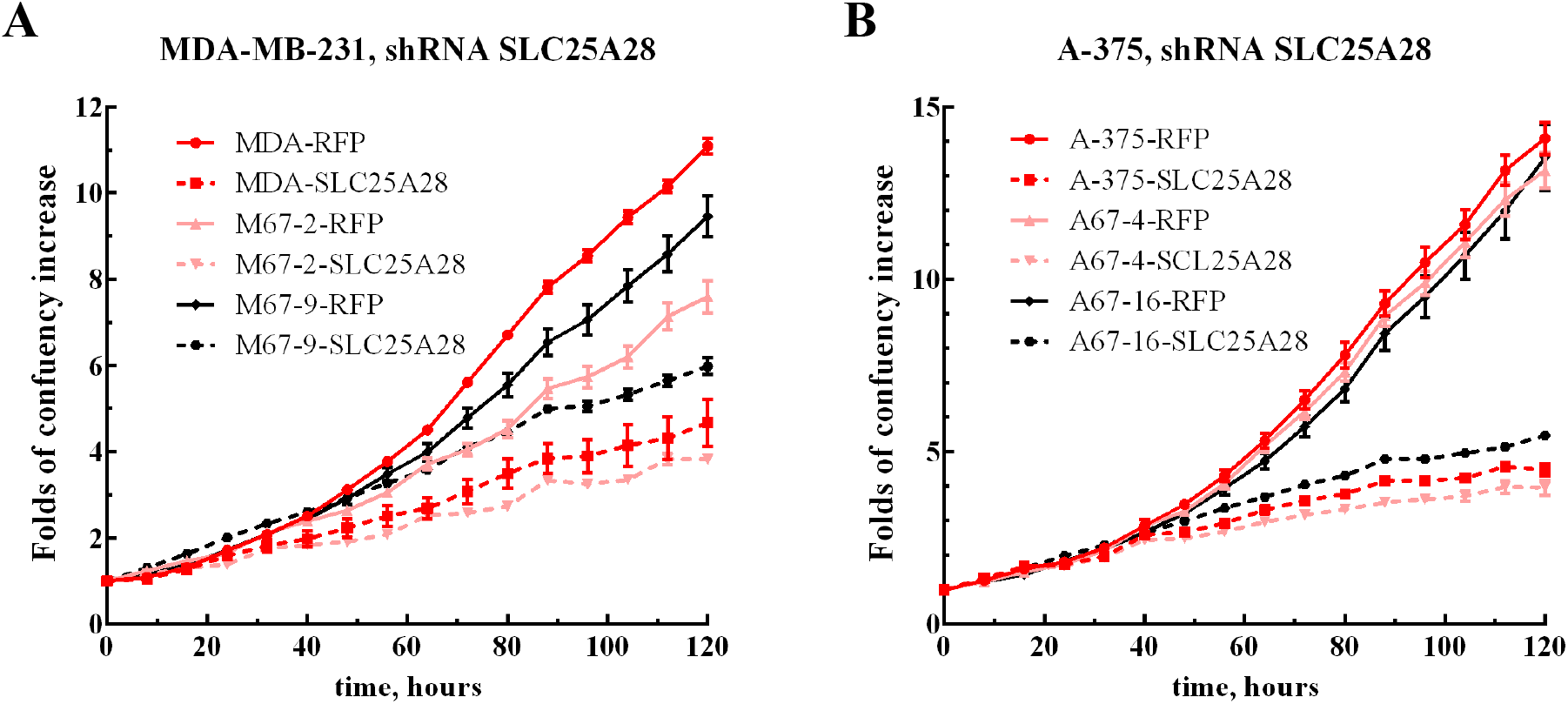
Growth curves of MDA-MB-231 (A) and A-375 (B) with high expression of MEMO1 (MDA-RFP and A-375-RFP, red solid lines), low expression of MEMO1 (M67-2-RFP and A67-4-RFP, pink solid lines) and no MEMO1 expression (M67-9-RFP and A67-16-RFP black solid lines), in comparison with these cell lines transduced with SLC25A28 shRNA (corresponding dashed lines).

**Figure S8.**
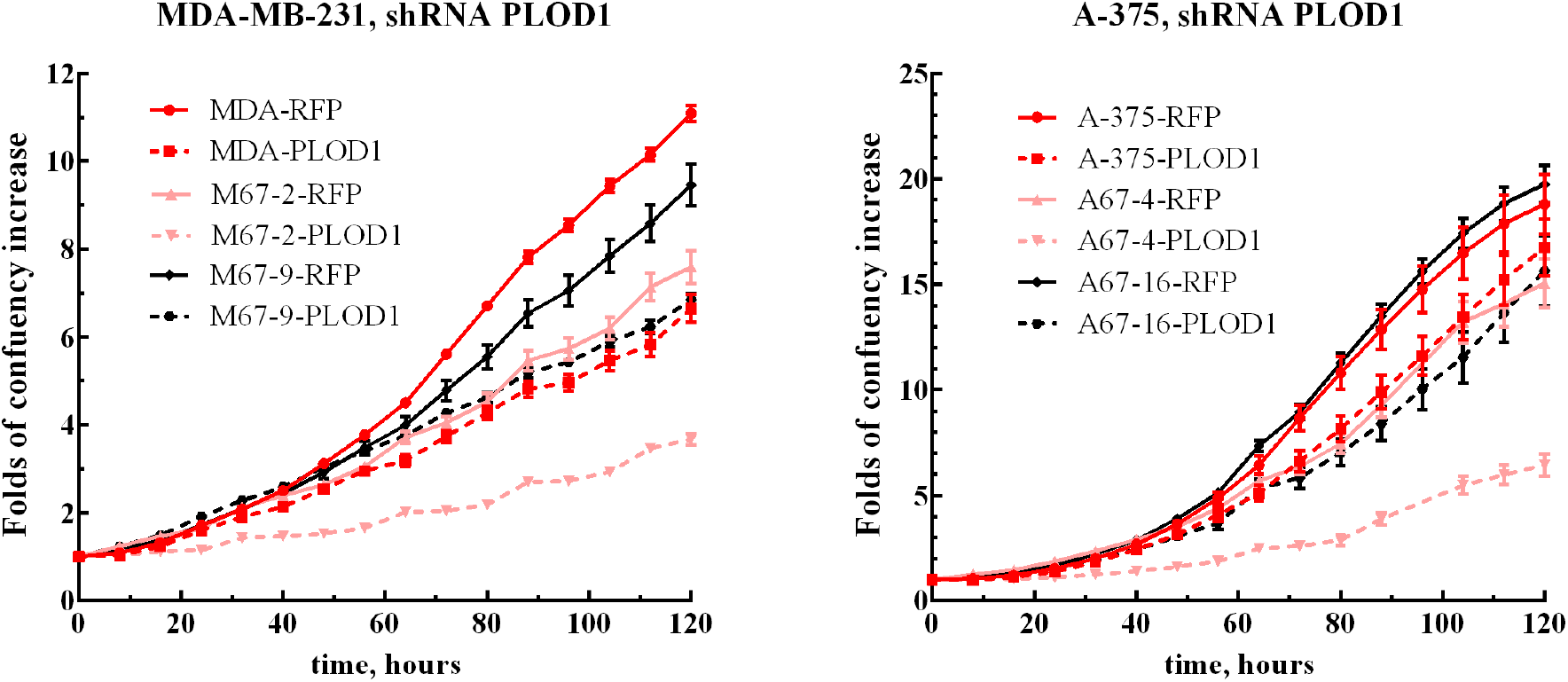
Growth curves of MDA-MB-231 and A-375 with high expression of MEMO1 (MDA-RFP and A-375-RFP, red solid lines), low expression of MEMO1 (M67-2-RFP and A67-4-RFP, pink solid lines) and no MEMO1 expression (M67-9-RFP and A67-16-RFP black solid lines), in comparison with these cell lines transduced with PLOD1 shRNA (corresponding dashed lines).

**Figure S9.**
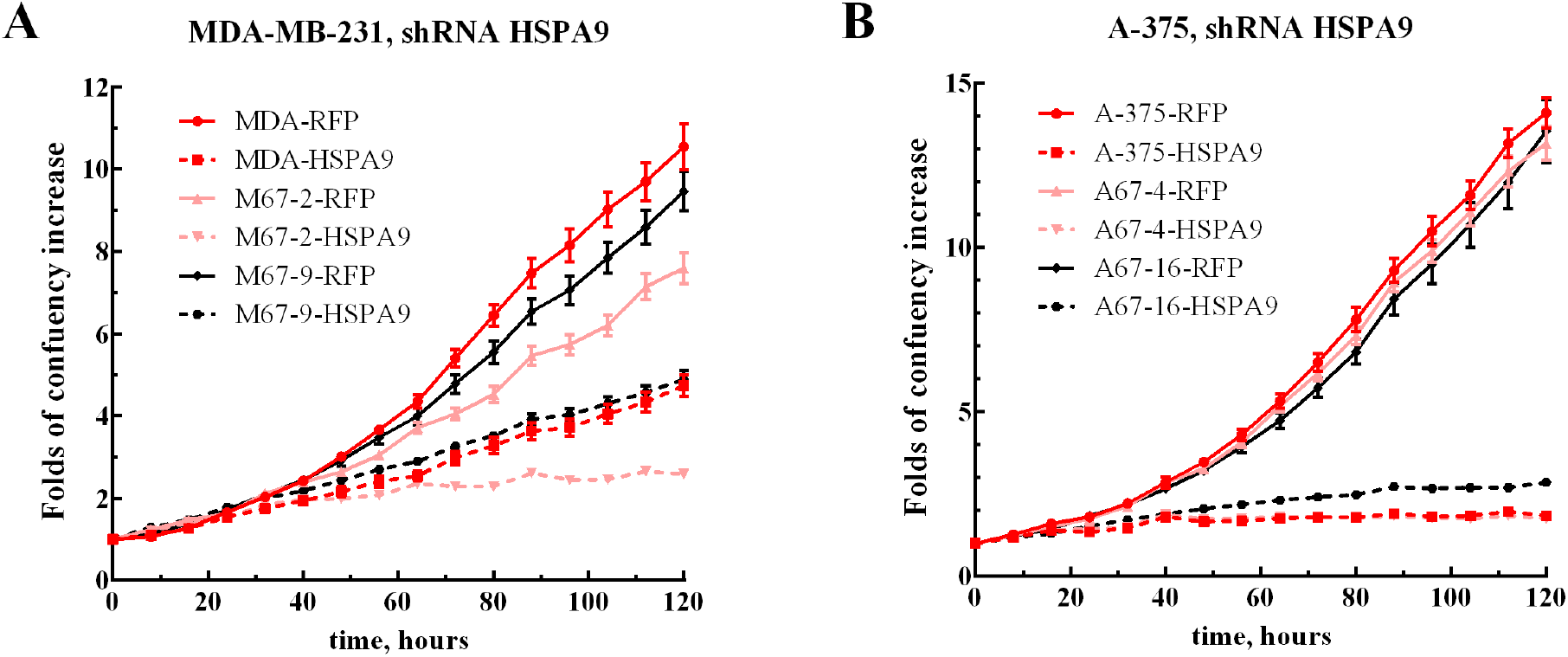
Growth curves of MDA-MB-231 (A) and A-375 (B) with high expression of MEMO1 (MDA-RFP and A-375-RFP, red solid lines), low expression of MEMO1 (M67-2-RFP and A67-4-RFP, pink solid lines) and no MEMO1 expression (M67-9-RFP and A67-16-RFP black solid lines), in comparison with these cell lines transduced with HSPA9 shRNA (corresponding dashed lines).

**Figure S10.**
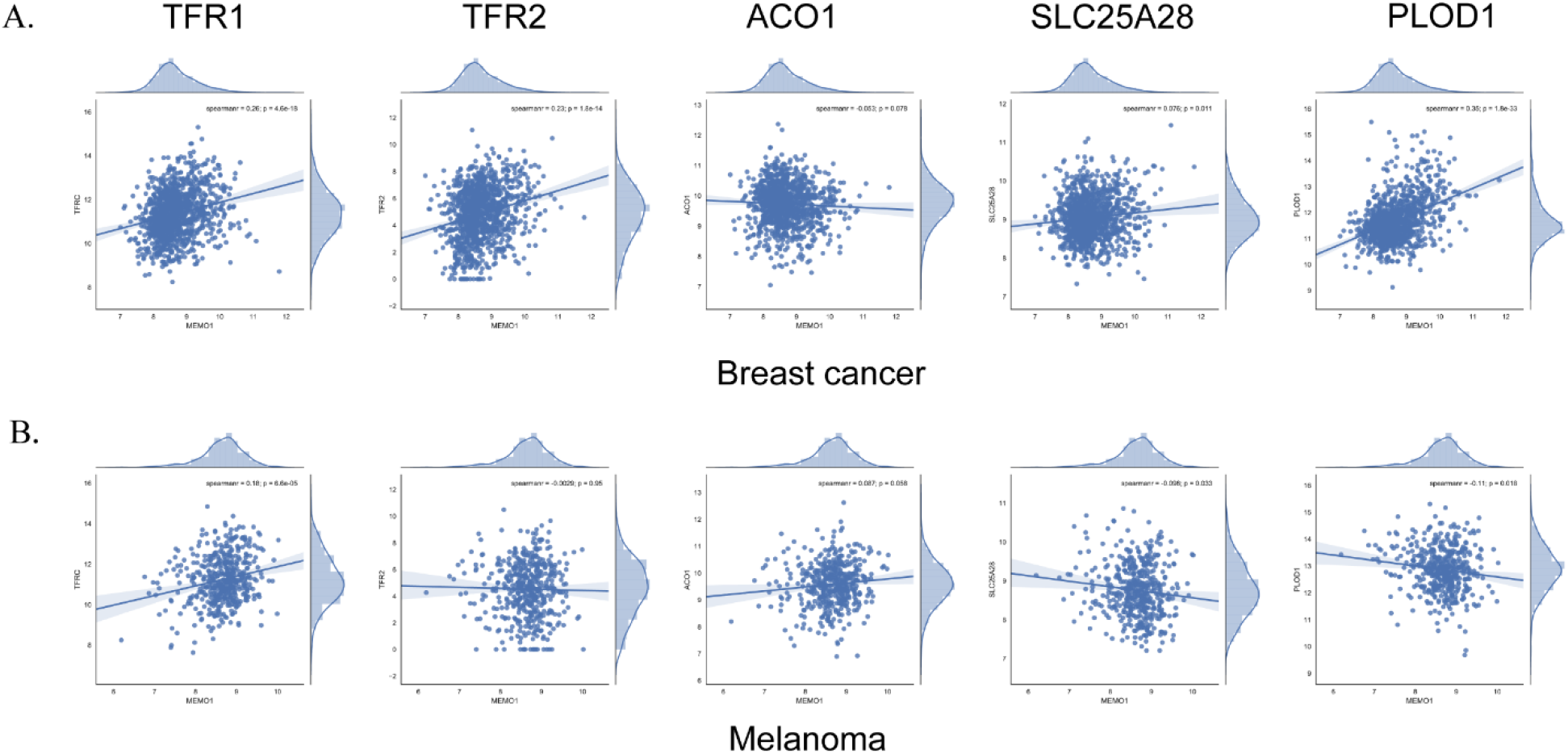
Correlation between the expression levels of MEMO1 and other iron related proteins in breast cancer (A) and melanoma (B) cell lines.

**Fig. S11.**
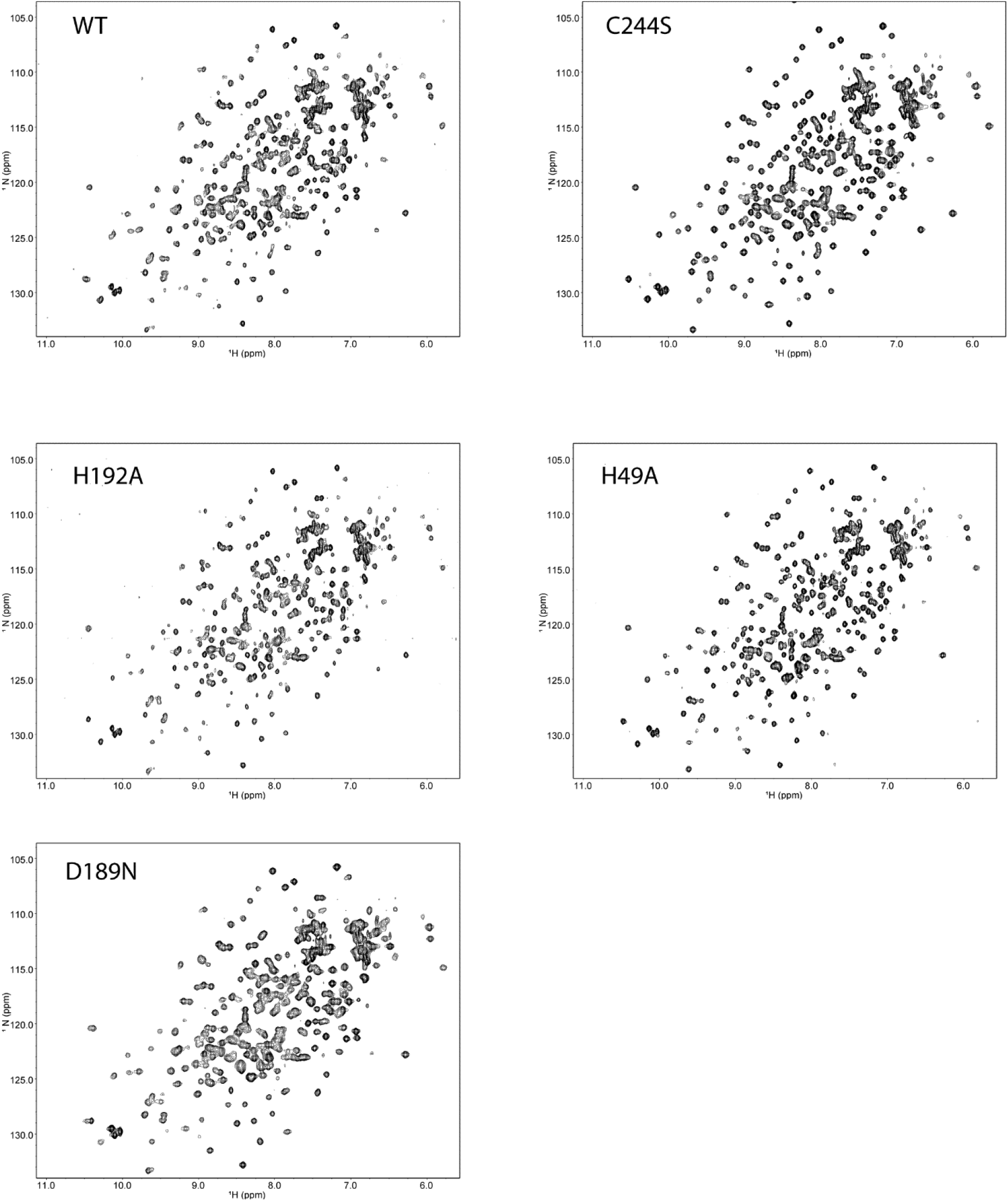
Fingerprint ^1^H,^15^N-TROSY spectra of the wild type MEMO1 and several metal binding site mutants recorded at 900 MHz. Pproteins were isotopically labeled with ^15^N by substituting ^15^NH_4_Cl for the natural abundance NH_4_Cl in the M9 medium used for protein expression. NMR samples contained 0.1 mM protein in 50 mM HEPES-Na, pH 7.4, 150 mM NaCl, 5 mM TCEP, 5% v/v D_2_O, and 0.25 mM 2,2-dimethyl-2-silapentane-5-sulfonate. The 2D ^1^H,^15^N-TROSY spectra were collected on a 900 MHz Bruker Avance III spectrometer equipped with a cryogenic triple-resonance probe at a sample temperature of 298 K.

**Fig. S12.**
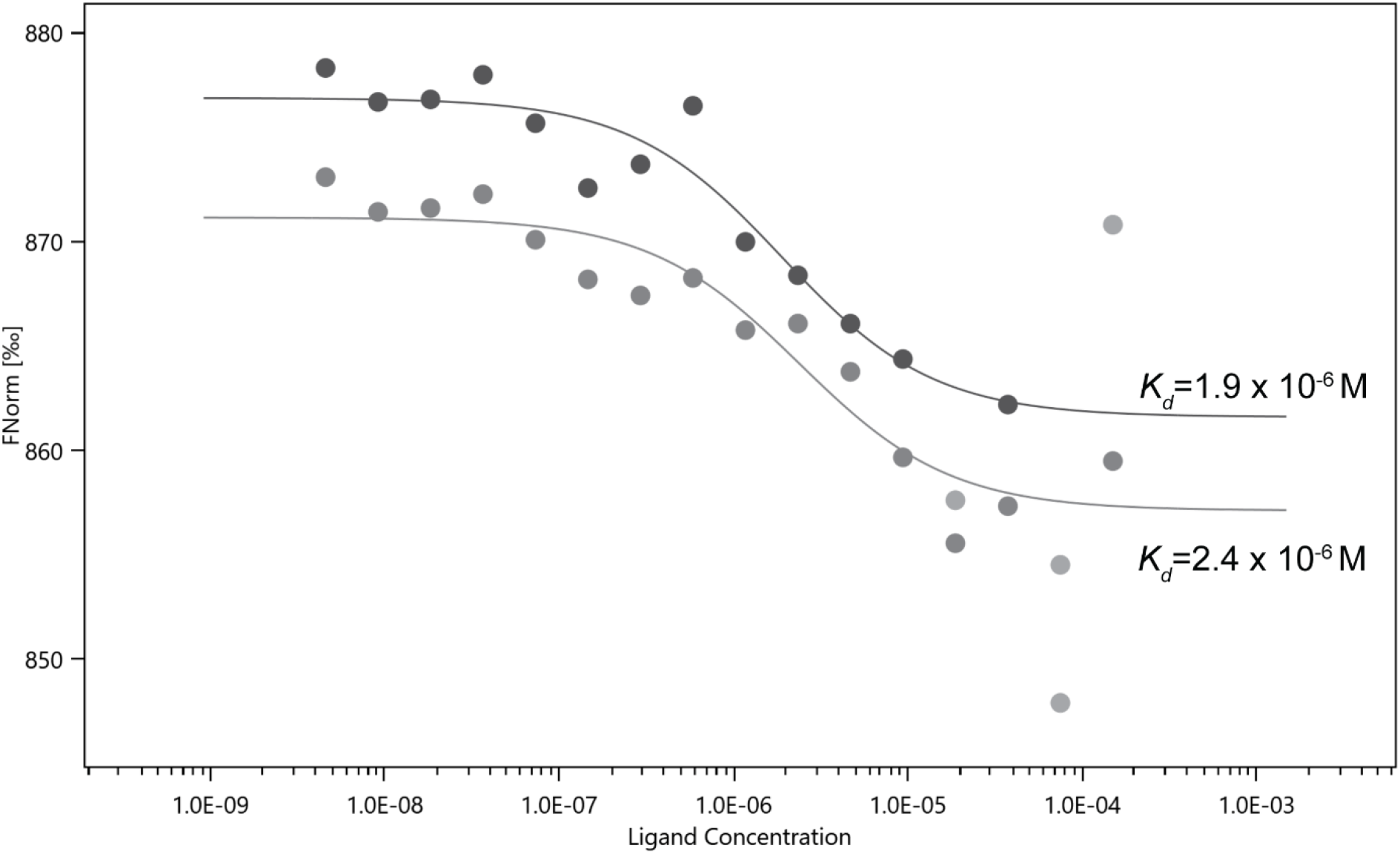
Iron binding to the wild type MEMO1 analyzed by microscale thermophoresis. Two independent experiments with calculated *K_d_* values are shown.

**Fig. S13.**
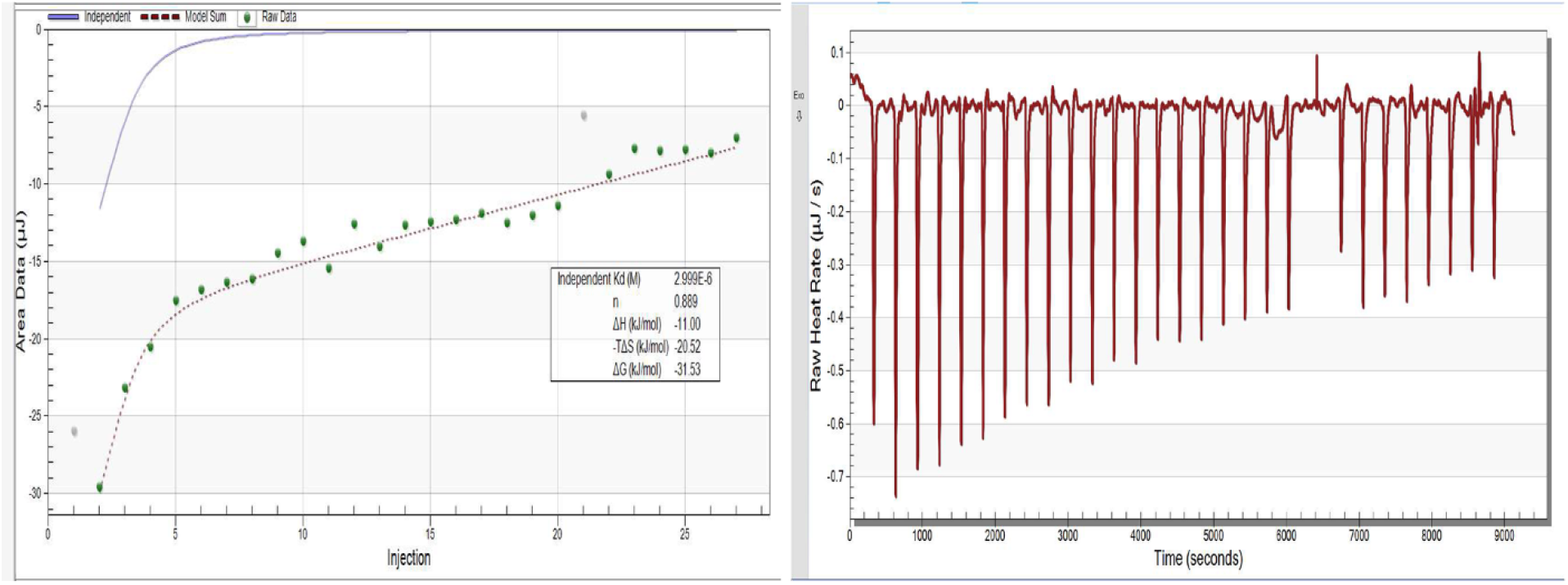
Copper binding to the wild type MEMO1 analyzed by isothermal titration calorimetry.

**Supplementary table S1.** Iron-related genes exhibiting GOF interactions with *MEMO1*.

| Gene | GO Slim | P-value | Distance low-high | Protein name | Database | Protein function |
| --- | --- | --- | --- | --- | --- | --- |
| *TFR2* | Ion regulation | 0.032 | 0.06 | Transferrin receptor 2 | CERES breast | Iron transport |
| *FTH1* | Immune System process | 0.037 | 0.19 | Ferritin H1 | DRIVE -RSA, pan-cancer | Cellular iron storage |
| *HSPA9* | Chaperone and Protein folding | 0.007 | 0.77 | Mitochondrial HSP70, Grp75, mortalin | Marcotte et al, pan-cancer | Iron-sulfur cluster biogenesis |
| *ACO1* | Metabolism | 0.019 | 0.16 | Apo-form: Iron-response protein (IRP); holo-form: Aconitase | DRIVE -Ataris, pan-cancer | Iron binding protein, aconitase/IRP |
| *ACO2* | Metabolism | 0.038 | 0.32 | Mitochondrial aconitase | CERES, pan-cancer | Fe<sub>4</sub>S<sub>4</sub>-cluster protein, TCA cycle |
|  |  | 0.032 | 0.66 |  | DRIVE -Ataris, pan-cancer |  |
| *HMOX1* | DDR pathway & NA metabolism | 0.043 | 0.51 | Heme oxygenase 1 | DRIVE -RSA, Breast | First enzyme of heme catabolism |
| *OGDH* | Mitochondrial organization | 0.036 | 0.62 | 2-oxoglutarate dehydrogenase | Achilles, pan-cancer | TCA cycle, regulates HIF1α |
|  |  | 0.002 | 0.22 |  | CERES, pan-cancer |  |
|  |  | 0.038 | 1.07 |  | Marcotte et al, pan-cancer |  |
| *SLC25A28* | Mitochondrial organization | 0.041 | 0.59 | Mitoferrin-2 | Marcotte et al, pan-cancer | Mitochondrial iron transport |
|  |  | 0.042 | 0.21 |  | CERES, Breast |  |
| *ISCU* | Mitochondrial organization | 0.023 | 1.78 | Iron-sulfur cluster assembly enzyme | Marcotte et al, pan-cancer | [Fe-S] cluster synthesis |
| *LIAS* | Mitochondrial organization | 0.0003 | 1.63 | Lipoic acid synthase | Marcotte et al, pan-cancer | Fe<sub>4</sub>S<sub>4</sub>-cluster containing protein, lipoate synthesis, regulates HIF1α |
|  |  | 0.009 | 1.74 |  | Marcotte et al, breast |  |
|  |  | 0.041 | 0.18 |  | CERES, pan-cancer |  |
| *PLOD1* | Protein modification | 0.045 | 0.41 | Procollagen-lysine,2-oxoglutarate 5-dioxygenase 1 | Marcotte, pan-cancer | Iron binding protein, regulates collagen synthesis, cross-linking, and deposition |
|  |  | 0.006 | 0.53 |  | Marcotte, breast |  |
|  |  | 0.042 | 0.07 |  | CERES, breast |  |
|  |  | 0.013 | 0.24 |  | Ataris, pan-cancer |  |
|  |  | 0.043 | 0.41 |  | DRIVE -Ataris, Breast |  |
|  |  | 0.010 | 0.21 |  | DRIVE -RSA, pan-cancer |  |

**Supplementary table S2.** Iron related genes exhibiting LOF interactions with *MEMO1*.

| Genes | GO Slim | P-value | Distance<br>low-high | Protein name | Database | Protein function |
| --- | --- | --- | --- | --- | --- | --- |
| *SLC11A2* | Apoptosis | 0.024 | -0.07 | DMT1 | CERES,<br>breast | Iron transport |
| *FTH1* | Immune System<br>process | 0.017 | -0.09 | Ferritin H1 | CERES<br>pan-cancer | Iron storage |
| *FXN* | Mitochondrial<br>organization | 0.032 | -0.79 | Frataxin | Marcotte<br>et al,<br>breast | [Fe-S] cluster biogenesis<br>in mitochondria |
| *FBXL5* | Miscellaneous | 0.038 | -0.57 | F-box/LRR-repeat protein 5 | Marcotte<br>et al, pan-cancer | regulates iron<br>homeostasis |
| *ISCA2* | Mitochondrial<br>organization | 0.038 | -0.39 | [Fe-S] cluster<br>assembly 2<br>homolog | Marcotte<br>et al, pan-cancer | [Fe-S] cluster biogenesis<br>in mitochondria |
| *NUBPL* | DDR pathways<br>& NA<br>metabolism | 0.013 | -0.69 | Iron-sulfur<br>protein NUBPL | Marcotte<br>et al, pan-cancer | [Fe-S] cluster biogenesis<br>in mitochondria |

**Supplementary table S3.** Genes involved in ferroptosis and exhibiting GOF or LOF interactions (highlighted in light blue) with *MEMO1*.

| Genes | GO Slim | P-value | Distance low-high | Protein name | Database | Protein function |
| --- | --- | --- | --- | --- | --- | --- |
| *TFR2* | Ion regulation | 0.032 | 0.06 | Transferrin receptor 2 | CERES, breast | Iron transport |
| *FTH1* | Immune System process | 0.037 | 0.19 | Ferritin H1 | DRIVE - RSA, pan-cancer | Cellular iron storage |
| *ACO1* | Metabolism | 0.019 | 0.16 | Apo-form: Iron-response protein (IRP); holo-form: Aconitase | DRIVE-Ataris, pan-cancer | Iron binding protein, aconitase/IRP |
| *VDAC1* | Ion Regulation | 0.0486 | 0.693 | Voltage-dependent iron channel 1 | Marcotte, pan-cancer | Transport of small hydrophilic molecules; regulation of cell volume and apoptosis |
| *VDAC3* | Ion regulation | 0.032 | 0.452 | Voltage-dependent iron channel 3 | Marcotte breast | Transport of small hydrophilic molecules; regulation of cell volume and apoptosis |
| *SLC25A28* | Mitochondrial organization | 0.041 | 0.59 | Mitoferrin-2 | Marcotte, pan-cancer | Mitochondrial iron transport |
|  |  | 0.042 | 0.21 |  | CERES, breast |  |
| *ACSL4* | Metabolism | 0.040 | 0.248 | Acyl-CoA Synthetase Long Chain Family Member 4) | Achilles pan-cancer | Lipid metabolism |
| *SIRT3* | unknown | 0.005 | 0.839 | Sirtuin-3 | Marcotte pan-cancer | NAD-dependent deacetylase |
|  |  | 0.042 | 0.554 |  | Marcotte breast |  |
|  |  | 0.009 | 0.539 |  | Achilles pan-cancer |  |
| *SCO2* | Mitochondrial organization | 0.001 | 0.149 | Sco2 | CERES pan-cancer | Mitochondrial copper chaperone |
| *ALOXE3* | metabolism | 0.030 | 0.063 | Hydroperoxide isomerase ALOXE3 | CERES pan-cancer | Non-Heme iron-containing lipoyxygenase |
| *PTGS2* | Intracellular Protein Traffic | 0.042 | 0.038 | prostaglandin-endoperoxide synthase 2 | CERES breast | Prostanoid biosynthesis |
| *FXN* | Mitochondrial organization | 0.032 | 0.787 | frataxin | Marcotte breast | [Fe-S] cluster biogenesis in mitochondria |
| *CS* | Metabolism | 0.048 | -0.236 | Citrate synthase | Ataris pan-cancer | TCA cycle |
| *ACSL4* | Metabolism | 0.042 | -0.418 | Acyl-CoA Synthetase Long Chain Family Member 4) | Marcotte breast | Lipid metabolism |
| *SCP2* | Cell cycle and Mitosis | 0.024 | -0.651 | sterol carrier protein 2 | Marcotte breast | Intracellular lipid circulation and metabolism |
| *GLS2* | Mitochondrial organization | 0.013 | 0.129 | Glutaminase 2 | CERES breast | Glutamine catabolism, mitochondrial respiration |
| *SCO2* | Mitochondrial organization | 0.042 | 0.744 | Sco2 | Marcotte breast | Mitochondrial copper chaperone |
|  |  | 0.022 | -1.32 |  | Achilles breast |  |
| *FH* | Mitochondrial organization | 0.014 | -0.328 | Fumarate hydratase | Atares pan-cancer | TCA cycle |
| *ALOX12* | Metabolism | 0.044 | 0.049 | arachidonate lipoxygenase | CERES pan-cancer | Lipid peroxidation |

**Supplementary Table S4.**
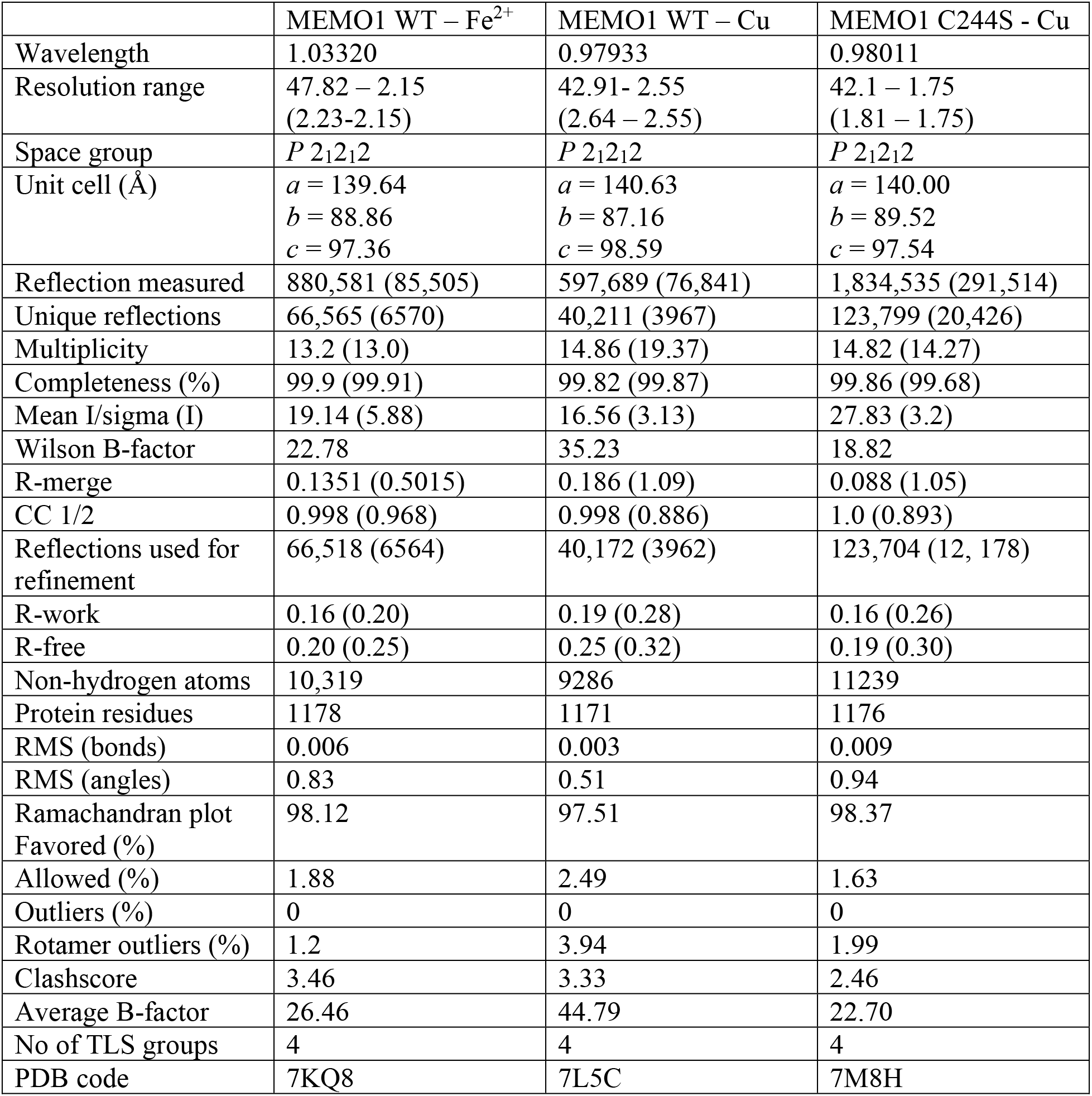
Structure determination statistics for MEMO1-metal complexes and the C244S mutant

|  | MEMO1 WT – Fe<sup>2+</sup> | MEMO1 WT – Cu | MEMO1 C244S - Cu |
| --- | --- | --- | --- |
| Wavelength | 1.03320 | 0.97933 | 0.98011 |
| Resolution range | 47.82 – 2.15<br>(2.23-2.15) | 42.91- 2.55<br>(2.64 – 2.55) | 42.1 – 1.75<br>(1.81 – 1.75) |
| Space group | *P* 2<sub>1</sub>2<sub>1</sub>2 | *P* 2<sub>1</sub>2<sub>1</sub>2 | *P* 2<sub>1</sub>2<sub>1</sub>2 |
| Unit cell (Å) | *a* = 139.64<br>*b* = 88.86<br>*c* = 97.36 | *a* = 140.63<br>*b* = 87.16<br>*c* = 98.59 | *a* = 140.00<br>*b* = 89.52<br>*c* = 97.54 |
| Reflection measured | 880,581 (85,505) | 597,689 (76,841) | 1,834,535 (291,514) |
| Unique reflections | 66,565 (6570) | 40,211 (3967) | 123,799 (20,426) |
| Multiplicity | 13.2 (13.0) | 14.86 (19.37) | 14.82 (14.27) |
| Completeness (%) | 99.9 (99.91) | 99.82 (99.87) | 99.86 (99.68) |
| Mean I/sigma (I) | 19.14 (5.88) | 16.56 (3.13) | 27.83 (3.2) |
| Wilson B-factor | 22.78 | 35.23 | 18.82 |
| R-merge | 0.1351 (0.5015) | 0.186 (1.09) | 0.088 (1.05) |
| CC 1/2 | 0.998 (0.968) | 0.998 (0.886) | 1.0 (0.893) |
| Reflections used for refinement | 66,518 (6564) | 40,172 (3962) | 123,704 (12, 178) |
| R-work | 0.16 (0.20) | 0.19 (0.28) | 0.16 (0.26) |
| R-free | 0.20 (0.25) | 0.25 (0.32) | 0.19 (0.30) |
| Non-hydrogen atoms | 10,319 | 9286 | 11239 |
| Protein residues | 1178 | 1171 | 1176 |
| RMS (bonds) | 0.006 | 0.003 | 0.009 |
| RMS (angles) | 0.83 | 0.51 | 0.94 |
| Ramachandran plot<br>Favored (%) | 98.12 | 97.51 | 98.37 |
| Allowed (%) | 1.88 | 2.49 | 1.63 |
| Outliers (%) | 0 | 0 | 0 |
| Rotamer outliers (%) | 1.2 | 3.94 | 1.99 |
| Clashscore | 3.46 | 3.33 | 2.46 |
| Average B-factor | 26.46 | 44.79 | 22.70 |
| No of TLS groups | 4 | 4 | 4 |
| PDB code | 7KQ8 | 7L5C | 7M8H |

